# Divergent somatic mutation patterns among human cerebellar neuron types

**DOI:** 10.1101/2025.09.29.679392

**Authors:** Marta Grońska-Pęski, Amoolya Srinivasa, Gilad D. Evrony

## Abstract

Neurons in the human brain accumulate somatic mutations with age. However, it is largely unknown how somatic mutation rates and patterns vary among the brain’s diverse types of neurons. Characterizing this variability is critical for elucidating the role of genome integrity in human brain function and disease. Moreover, the significant physiological differences among the brain’s cell types provides an opportunity to learn more general underlying factors that determine mutation rates and patterns. Here, we utilized high-fidelity duplex DNA sequencing to profile somatic mutation processes across the lifespan in the two major cell types of the human cerebellum, Purkinje neurons and granule neurons, which have dramatically different sizes, functions, and physiologies. Surprisingly, these cell types exhibited similar rates of substitution mutations, including similar rates of signature SBS5 that is responsible for most mutations in the body yet whose mechanism remains unknown. However, we identified differences in Purkinje and granule neurons’ patterns of substitutions and in their rates and patterns of insertions and deletions, with transcription playing a key role in mediating these differences. Our work indicates that different types of neurons in the brain can differ in their aging-related somatic mutation processes. Our results further suggest that key features that distinguish Purkinje neurons from granule neurons, such as cell size, metabolic rates, and neuronal firing rates, are unlikely to be intrinsic determinants of the total substitution mutation rate and of signature SBS5, which is the most prevalent aging-related mutational process.

## Main

Every cell in the body accumulates somatic mutations with age^1–7^, but how somatic mutation rates and patterns vary among different cell types remains largely unexplored. In the human brain, aging-related somatic mutations have only been profiled genome-wide in non-specific populations of cortical and hippocampal neurons^2,4,6^ and in large cortical neurons^5^, as well as in oligodendrocytes^4^. Characterizing whether and how genome-wide somatic mutation rates and patterns vary among the brain’s diverse cell types, especially among specific neuronal cell types, is critical to understanding healthy brain aging as well as aging-related brain diseases, which often preferentially affect specific types of neurons^7,8^. For example, neurodegeneration in Huntington’s disease has recently been attributed to aging-related somatic DNA repeat expansion in the *HTT* gene that occurs only in specific types of neurons such as striatal projection neurons^7,8^.

The human cerebellum contains approximately 80% of the brain’s neurons^9,10^. Two of the major types of neurons in the cerebellum, Purkinje neurons (PNs) and granule neurons (GNs), have important roles in motor control, coordination, and sensory processing^11^. While PNs and GNs are located in close proximity and communicate directly with each other (GN axons synapse extensively with PN dendrites), these two neuronal cell types have remarkably different sizes, morphologies, metabolic activities, and electrical activities^9,12–18^. For example, PNs and GNs exhibit perhaps the most extreme size difference among functionally interacting cells in the body. Among all neuronal cell types in the human brain, PNs have the second largest cell body diameters (35 to 65 µm) compared to GNs that have the smallest cell body diameters (∼5 µm diameter), and PNs have an estimated 200 times greater cell body volume (excluding their axonal and dendritic trees) than GNs^13,15,19,20^. While PNs and GNs in animal models both have bursts of firing with rates up to ∼250 Hz, their basal firing rates are markedly different: between 30 to 150 Hz in PNs compared to < 3 Hz in GNs^17,18,21–25^. A computational analysis integrating these and other aspects of PN and GN physiology in rodents estimated that PNs consume ∼60 times more ATP per second than GNs on a per-cell basis^25^.

These and other physiological differences likely mediate the differential susceptibility of PNs and GNs in disease states^26–28^. For example, PNs are preferentially vulnerable and lost in many conditions, including after ischemia, in diverse genetic diseases (e.g., SCA1, SCA6, and SCA14 spinocerebellar ataxias; Friedreich’s ataxia; NPC1 lysosomal storage disease; PEX7 peroxisome biogenesis disorder; and ataxia with vitamin E deficiency), and due to some toxins (e.g., ethanol)^26–35^. In contrast, GNs are preferentially affected in other genetic diseases (e.g., *TUBB4A* leukoencephalopathy) and by other toxins (e.g., methylmercury)^28,36^. Additionally, some conditions involving the cerebellum affect both PNs and GNs, such as SCA3 and SCA7 spinocerebellar ataxias, ataxia telangiectasia, and Alzheimer’s disease—though when both cell types are affected, it can be uncertain whether loss of one cell type is secondary to loss of the other^27,32,37–39^.

However, it is unknown whether these striking physiological differences between PNs and GNs—while sharing an anatomic niche—are associated with differences in their ability to maintain genomic integrity. Determining this for these two significantly different cell types may, in turn, help inform more generally whether cell size and cellular metabolism are major contributors to somatic mutation rates. Here, we developed an approach to profile human cerebellar PNs and GNs by high-fidelity duplex DNA sequencing, which revealed the landscape of somatic mutation in these neuronal cell types across the lifespan.

### Isolating cerebellar Purkinje and granule neurons

We obtained postmortem frozen human cerebellum from 21 neurotypical females across the lifespan: children (2-19 years old, yo; N=4), young adults (20-39 yo; N=4), middle-aged adults (40-59 yo; N=4), old adults (60-79 yo; N=4), and older adults (≥80 yo; N=5) (**Supplementary Table 1**). The cell bodies of PNs and GNs that contain their nuclear genomes are located in anatomically distinct layers^19,40^, which we reasoned could allow us to isolate PNs and GNs by microdissection of histologic sections. Specifically, PN cell bodies have readily identifiable morphology and are sparsely arranged along the approximately one-cell-thick Purkinje cell layer, while nearly all the cells in the adjacent thicker granule cell layer are GNs^9,41^.

We therefore sectioned the frozen cerebellar tissues, and after staining with hematoxylin and eosin, we dissected out PN cell bodies one at a time using laser-capture microdissection (LCM) to obtain an average of ≍ 900 PNs per individual and a total of 19,255 PNs across all individuals (**Figs. 1a,b, Supplementary Table 2, and Methods**). To validate the cell type identity of LCM-isolated PNs, we performed Smart-seq3^42^ bulk RNA-sequencing (RNA-seq) of PNs isolated from three individuals (**Methods**). PN samples showed high expression of canonical PN markers, with negligible expression of other cell type markers (**Fig 1c. and Supplementary Table 3**), confirming that our LCM procedure obtains nearly pure populations of PNs.

**Figure 1.**
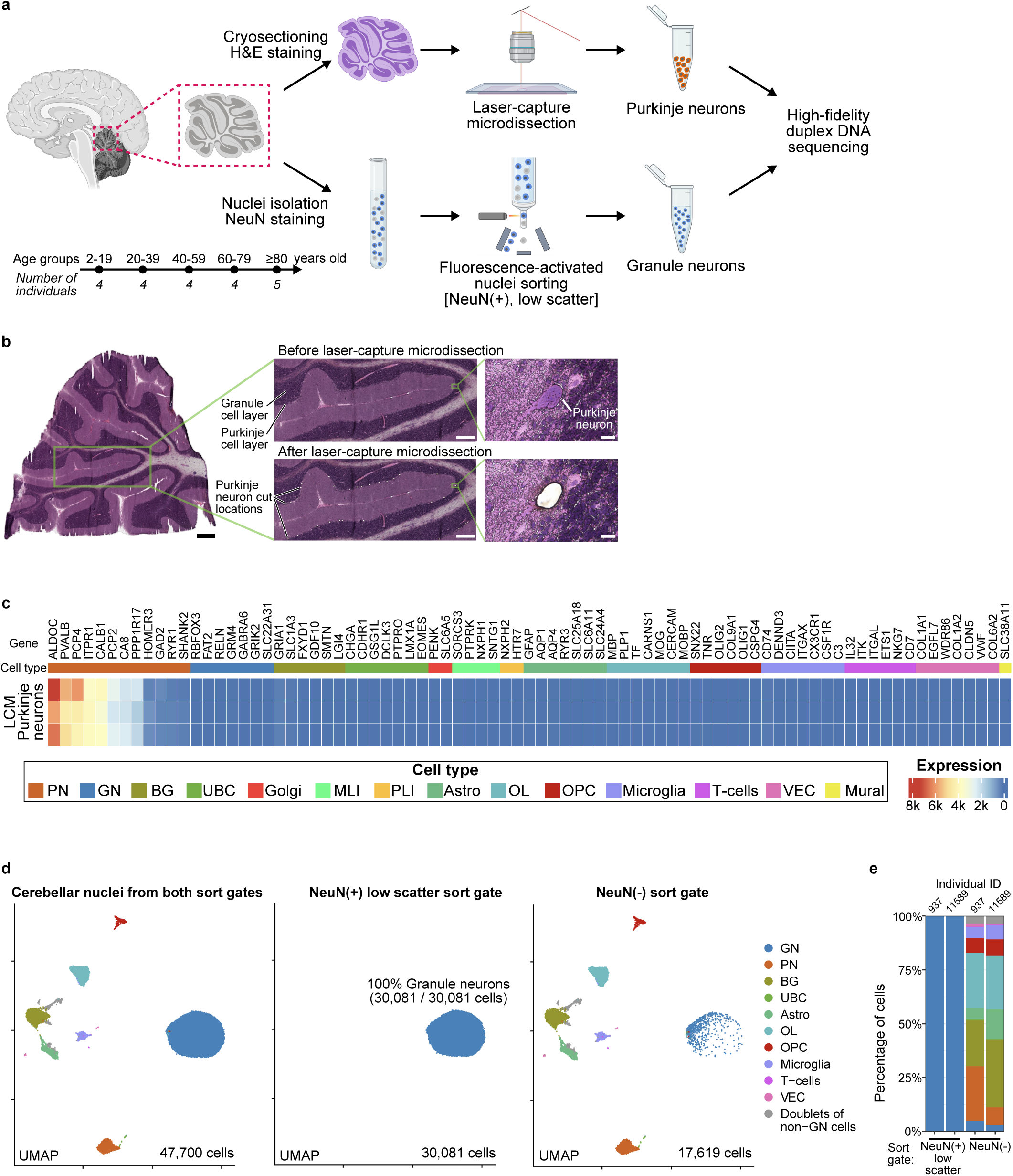
Purification of Purkinje neurons and granule neurons. **a**, Schematic of profiled samples and method. H&E, hematoxylin and eosin. Created in BioRender. Evrony, G. (2025). **b**, Representative laser-capture microdissection of Purkinje neurons. Scale bars: 1 mm, left panel; 500 µM, middle panels; 20 µM, right panels. **c,** Normalized unique molecular index (UMI) exon-aligned read counts obtained by Smart-seq3 bulk RNA-seq of three laser-capture microdissections of Purkinje neurons (top to bottom, individuals 937, 5609, 11589) for marker genes of major cerebellar cell types. See **Supplementary Table 3** for data of all genes. k, × 1000. **d,** snRNA-seq UMAP plots of fluorescence-activated nuclei sorting from cerebellum. The UMAP plots combine the nuclei of both individuals profiled in this experiment (937 and 11589). The middle plot of nuclei sorted using the NeuN(+) low scatter sort gate corresponds to the sort gate used for granule neuron mutation profiling. ‘Doublets of non-GN cells’ are several clusters containing doublets of two different non-granule neuron cell types (**Methods**). See **Extended Data Figs. 2a-c** for details of sort gates and expression of marker genes of annotated cell types. **e,** The cell type fractions of nuclei sorted by each sort gate, using the same color legend as in Fig. 1d. See **Supplementary Table 4** for nuclei counts of each cell type. **c,e,** GN: granule neurons; PN: Purkinje neurons; BG: Bergmann glia; UBC: unipolar brush cells; Golgi: Golgi neurons; MLI: molecular layer interneurons; PLI: Purkinje layer interneurons; Astro: astrocytes; OL: oligodendrocytes; OPC: oligodendrocyte progenitor cells; VEC: vascular endothelial cells; Mural: mural cells.

Since GNs are too small to isolate one at a time using LCM, we evaluated whether we could purify GNs using LCM by cutting from the granule cell layer of each of three individuals approximately 250 – 300 circles, each of which is ≍ 20 µm in diameter containing ≍ 10 GN cell bodies. However, bulk RNA-seq of these samples showed co-purification of GNs with astrocytes and oligodendrocytes (**Extended Data Fig. 1 and Supplementary Table 3**). We therefore developed an alternative approach for isolating pure populations of GNs using fluorescence-activated nuclei sorting (FANS). We stained cerebellar nuclei with anti-NeuN antibody and then sorted from the predominant NeuN(+) population that had a low scatter signal, as GNs have the smallest nuclei among all cerebellar neurons^16^ (**Extended Data Figs. 2a,b and Methods**). To validate the cell type identity of FANS-sorted GNs, we performed single-nucleus RNA-seq (snRNA-seq) of NeuN(+) low-scatter population and of the NeuN(-) population from each of two individuals (**Methods**). The NeuN(+) low scatter population showed remarkable purity of GNs: 30,081 / 30,081 (100%) of sorted nuclei in this population were classified as granule neurons (**Fig. 1d, Extended Data Fig. 2c, and Supplementary Table 4**), confirming that our FANS procedure obtains pure populations of GNs.

### Mutation rates

We proceeded to profile somatic mutations in PNs and GNs using the NanoSeq duplex DNA sequencing method^2^ that we further optimized for low DNA input (**Methods**). Duplex DNA sequencing dramatically reduces sequencing error rates via a consensus of reads from both strands of each DNA molecule^43^, and NanoSeq is a recently developed type of duplex sequencing that eliminates duplex artifacts that can arise during library preparation, achieving single-molecule fidelity in detecting somatic mutations without any discernible artifacts^2^. We obtained a total of 13.9 terabases of sequencing data (average 331 gigabases per sample) for somatic mutation analysis (**Supplementary Table 2**). After duplex consensus collapsing and filtering, this yielded an average of 3.8 × 10^9^ interrogated base pairs per sample (∼1.2 haploid genome equivalents) (**Supplementary Table 5**). Germline variants were filtered using additional sequencing data that we obtained for each individual from either bulk kidney, liver, cerebral cortex, or cerebellum (**Supplementary Table 2**). We performed a fingerprinting analysis using germline variants to confirm that all the sequencing samples of each individual derived from the same individual (**Extended Data Fig. 3a and Methods**). Across all samples, we detected a total of 29,721 somatic substitution mutations and 4,270 somatic insertion/deletion (indel) mutations (**Supplementary Tables 5 and 6**).

Both PNs and GNs exhibited an age-related increase in the burden of somatic substitution and indel mutations (**Figs. 2a,b and Supplementary Table 7**). PNs and GNs accrued 4.2 × 10^-9^ (95% confidence interval [CI]: 4.0 – 4.4 × 10^-9^) and 4.0 × 10^-9^ (95% CI: 3.8 – 4.3 × 10^-9^) substitutions per base pair (bp) per year, respectively, and 7.4 × 10^-10^ (95% CI: 6.9 – 7.9 × 10^-10^) and 3.6 × 10^-10^ (95% CI: 3.2 – 4.0 × 10^-10^) indels per bp per year, respectively (**Figs. 2a,b**). Per PN and GN cell, this corresponds to 26.1 and 24.9 substitutions per year, respectively, and 4.6 and 2.2 indels per year, respectively. Surprisingly, despite the significant differences in PN and GN cell size and physiology, their substitution mutation rates were not significantly different (rate ratio 1.05; 95% CI: 0.98 – 1.11; *P* = 0.14; negative binomial model with a two-sided Wald test; **Methods**) (**Fig. 2a and Supplementary Table 8**). In contrast, the indel mutation rate was approximately twice as high in PNs than in GNs (rate ratio 2.06; 95% CI: 1.84 – 2.31; *P* < 10^-15^) (**Fig. 2b and Supplementary Table 8**). The substitution rate of PNs was only slightly higher than that of cerebral cortex neurons profiled in a prior duplex sequencing study^2^ (rate ratio 1.11; 95% CI: 1.01 – 1.22; *P* = 0.03), while GN and cortical neuron substitution rates did not significantly differ (*P* = 0.35) (**Extended Data Fig. 3b**). Cortical neuron indel mutation rates were intermediate between PN and GN rates (**Extended Data Fig. 3c**).

**Figure 2.**
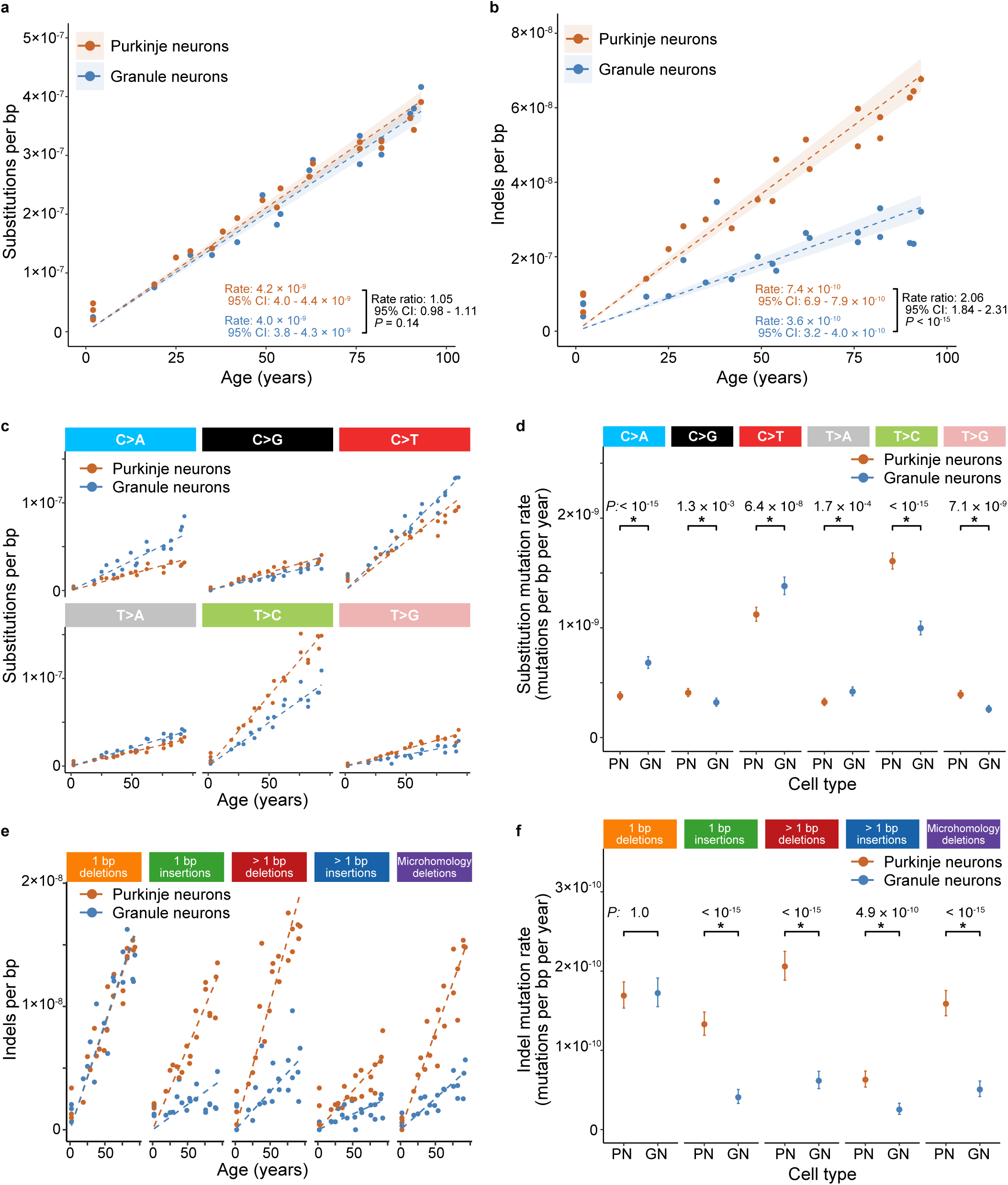
Substitution and indel mutation rates. **a,b,** Substitution (a) and indel (b) mutation burdens versus age. Rates are in units of mutations/[bp · year]. CI, confidence interval. **c-f,** Substitution (c) and indel (e) mutation burdens versus age for different mutation types, and comparisons between PN and GN substitution (d) and indel (f) mutation rates for different mutation types. **a,b,c,e,** Dashed lines (a,b,c,e) show predictions of the regression model and shaded regions (a,b) show 95% confidence bands (**Methods**). **d,f,** PN versus GN comparisons show *P*-values (asterisks mark significant differences) and error bars show 95% Cis (**Methods**). See **Supplementary Tables 7,8** for further details of rates and comparison statistics.

Although total PN and GN substitution rates did not differ, when we compared rates separately for each of the six possible substitutions (i.e., C>A, C>G, C>T, T>A, T>C, T>G), we identified significant differences for each. Specifically, PNs had higher C>G, T>C, and T>G mutation rates, while GNs had higher C>A, C>T, and T>A mutation rates (**Figs. 2c,d and Supplementary Tables 7,8**). Interestingly, these difference in rates of specific substitution types did not yield a difference in total PN and GN substitution rates, because the sum of the differences in rates of substitution types that were higher for PNs was nearly the same as the sum of the differences in rates of substitution types that were higher in GNs (**Supplementary Table 7**). Moreover, when we compared rates separately for different types of indels (1 bp deletions, 1 bp insertions, > 1 bp deletions, > 1 bp insertions, and microhomology deletions), we observed significant differences: PNs had higher rates than GNs for all indel types except 1 bp deletions (**Figs. 2e,f and Supplementary Tables 7,8**). Overall, these results indicate that PNs and GNs, despite having similar substitution rates as well as total mutation rates (as indels account for only a minority of mutations), exhibit divergent age-related mutational patterns.

### Mutational signatures

We further explored the differences in PN and GN mutational patterns by plotting mutational spectra and performing mutational signature analyses for substitutions across the full spectrum of 96-trinucleotide contexts and for indels across 83 types of indels^44^ (**Supplementary Table 9**).

The spectrum of all PN substitutions combined was largely similar to the spectrum of all GN substitutions combined (cosine similarity 0.89), but with notable differences—for example, a greater fraction of PN substitutions are ATN>ACN and a greater fraction of GN substitutions are C>A and NCG>NTG (**Fig. 3a**). Next, we performed mutational signature analysis of substitutions, excluding samples from the three two-year-old individuals as their samples’ mutation counts were too low to yield reliable mutational spectra. Signature extraction yielded signatures resembling COSMIC SBS1, SBS5, and SBS16, though with low cosine similarities of the extracted SBS1-like and SBS16-like signatures to their corresponding COSMIC catalogue signatures^44^ (**Methods**) due to the challenge in extracting the low level of SBS1 from small sample cohorts^2,45^ and the challenge of separating SBS5 from SBS16^46^. Therefore, to obtain more accurate exposure estimates for these signatures, we fit our data to SBS1, SBS5, and SBS16 from the COSMIC catalogue. This analysis showed that SBS1 mutation rates and average exposures (i.e., the fraction of substitutions due to the signature) were lower in PNs (95% CI of PN/GN rates: 0.39 – 0.60; average exposures: 0.06 and 0.13 in PNs and GNs, respectively), SBS5 mutation rates and average exposures did not differ significantly (95% CI of PN/GN rates: 0.94 – 1.08; average exposures: 0.80 and 0.82 in PNs and GNs, respectively), and SBS16 mutation rates and average exposures were higher in PNs (95% CI of PN/GN rates: 2.12 – 3.45; average exposures: 0.14 and 0.05 in PNs and GNs, respectively) (**Figs. 3b,c, Extended Data Fig. 4, and Supplementary Table 10**). Recent studies have closely linked SBS1 to cell division^47^, yet in both PNs and GNs, there was a low but non-zero rate of SBS1 (2.7 × 10^-10^ and 5.5 × 10^-10^ mutations/[bp · yr], respectively). This was also observed in a recent study of cortical neurons^46^. This finding may either be due to imperfect estimation of signature exposures or it may indicate that SBS1 can occur at low rates post-mitotically. The higher burden of SBS1 in GNs than PNs may also reflect the more numerous cell divisions during development and the early postnatal period that create the large population of GNs^48^. The higher rate in PNs of SBS16, which prior studies have closely linked to transcription^49,50^, is consistent with PNs having greater transcriptional activity and transcription-related mutagenesis than GNs.

**Figure 3.**
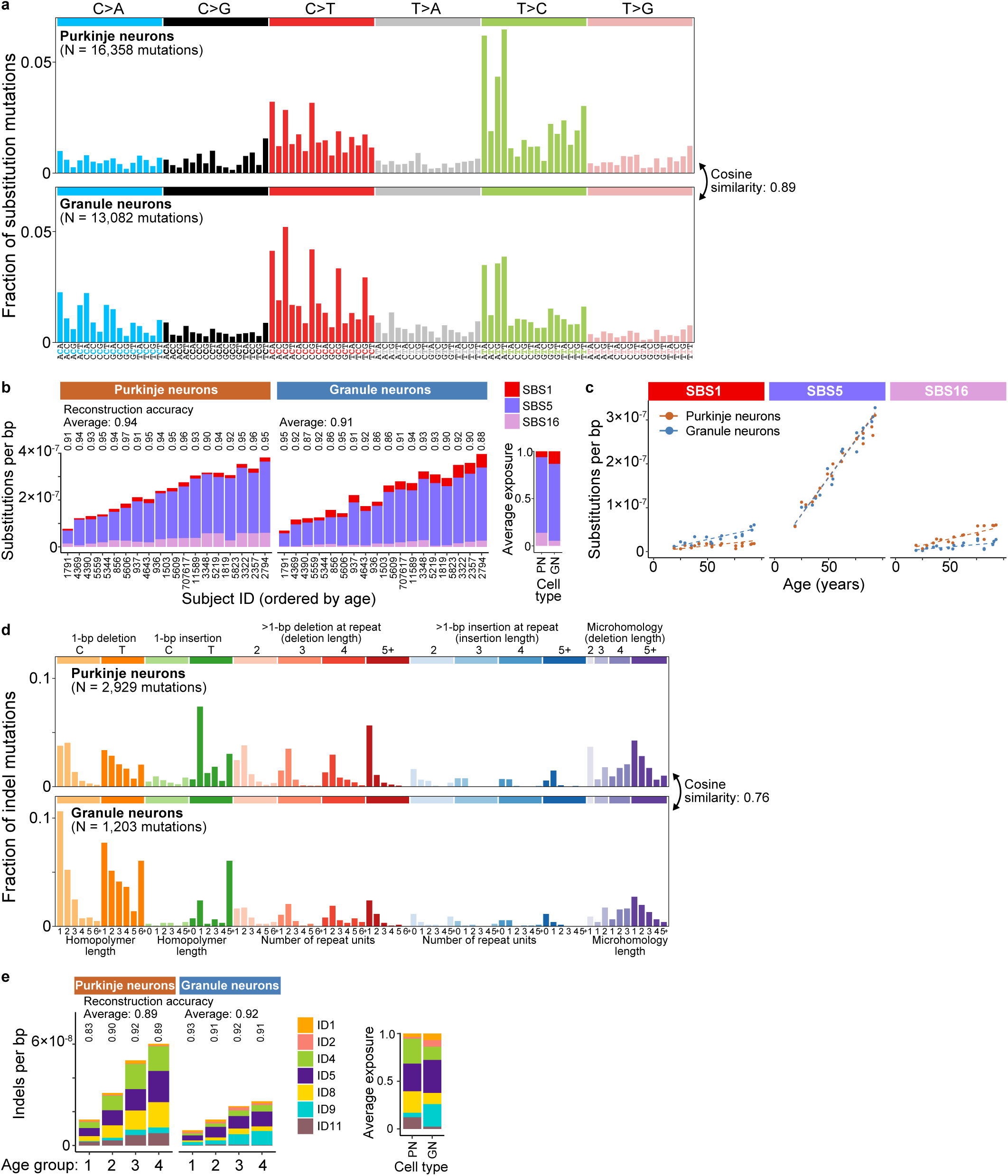
Mutational spectra and signatures. **a,** Mutational spectra of unique substitutions (i.e., counting only once those mutations that are detected more than once within a sample) combined across all samples of each cell type, corrected for trinucleotide context opportunities^2^ (i.e., the distribution of trinucleotides observed in interrogated base pairs relative to the distribution of trinucleotides in the genome). **b,** Burdens of substitution (SBS) signatures in each sample, excluding samples from two-year-old individuals due to low mutation counts that make signature analysis unreliable. Reconstruction accuracy is annotated above each sample, as well as the average reconstruction accuracy across samples of each cell type. On the right is the exposure (i.e., fraction of mutations) of each signature averaged across samples of each cell type. **c,** Substitution mutation burdens versus age for each signature. Dashed lines show predictions of the regression model (**Methods**), and see **Extended Data Fig. 4** for comparisons of PN versus GN rates. **d,** Mutational spectra of unique indels combined across all samples of each cell type. **e,** Burdens of indel (ID) signatures in each sample, after grouping samples by age (Group 1: 2-29 yo (6 individuals); Group 2: 30-53 yo (5 individuals); Group 3: 54-79 yo (5 individuals), Group 4: ≥ 80 yo; **Methods**). Reconstruction accuracies are annotated as in (b). On the right is the exposure (i.e., fraction of mutations) of each signature averaged across age groups of each cell type. **b,e,** See **Supplementary Table 10** for exposure estimates. PN, Purkinje neurons; GN, granule neurons.

Overall, our data show that PNs and GNs have similar total substitution rates primarily because they have similar rates of SBS5, which accounts for most of their substitution mutations, while SBS1 and SBS16 have nearly reciprocal rate differences. These results suggest more generally that the major properties by which PNs and GNs differ, such as cell size, metabolic rates, and neuronal firing rates, may not be intrinsic determinants of the rate of SBS5, which gives rise to most mutations in the body.

The spectra of PN and GN indels were significantly more divergent (cosine similarity 0.76) than substitution spectra, suggesting that indels arise from more cell type-specific mutational mechanisms (**Fig. 3d**). Notable differences included PNs having a greater fraction of indels that are 1 bp insertions of T adjacent to single (non-homopolymer) T bases and a greater fraction of indels that are > 1 bp deletions, as well as GNs having a greater fraction of indels that are 1 bp deletions and a greater fraction of indels that are 1 bp insertions of T in long (≥ 5 bp) homopolymers (**Fig. 3d**). Due to the significantly lower number of indels relative to substitutions, we performed indel signature analysis after pooling samples into four age groups (**Methods**). Since the low total number of indels also led de novo signature extraction to yield signatures with low similarity to the COSMIC catalogue, we fit our data to a set of known COSMIC signatures (ID1, ID2, ID4, ID5, ID8, ID9, and ID11) that are either prevalent across tissues or that were observed in a prior study of somatic mutations in neurons^44,46^ (**Methods**). This analysis identified that the most active signatures in PNs are ID4, ID5, ID8, and ID11 with average exposures of 0.26, 0.29, 0.23, and 0.12, respectively, whereas the most active signatures in GNs are ID4, ID5, and ID9 with average exposures of 0.14, 0.35, and 0.24, respectively (**Fig. 3e and Supplementary Table 10**). Both ID4 and ID11 have been previously associated with transcription^49,51^, so their higher exposure in PNs is again consistent with PNs having greater transcription-related mutagenesis than GNs. ID8 may arise from non-homologous-end-joining repair of DNA double-strand breaks^44^, suggesting that such DNA breaks occur more often in PNs than in GNs.

### Associations with transcription and chromatin accessibility

To further investigate the possible mechanisms underlying the differences in PN and GN somatic mutation rates and patterns, we profiled somatic mutations in genomic regions stratified by transcriptional activity, by transcribed versus untranscribed strand, and by chromatin accessibility.

For analyses stratified by transcriptional activity, we used our above snRNA-seq data of PNs and GNs to divide the genome into six sets of regions for each cell type: intergenic regions, regions of genes that are not expressed (Q0), and regions of genes with non-zero expression binned into four quartiles of expression levels (Q1-Q4; Q1, lowest expression, Q4, highest expression) (**Supplementary Table 11, Extended Data Figs. 5a,b, and Methods**). We then quantified the somatic mutation rates of each sample within each of its respective cell type’s six sets of regions, normalizing for the number of interrogated base pairs of each region set in each sample.

Surprisingly, although the total substitution mutation rates of PNs and GNs are not significantly different (**Fig. 2a**), this analysis revealed that PNs have higher mutation rates than GNs in genomic regions with the highest quartiles of gene expression (95% CI of PN/GN rates: Q2 = 1.25 – 1.70, Q3 = 1.48 – 1.98, Q4 = 1.69 – 2.15) while GNs have a higher mutation rate than PNs in intergenic regions (95% CI of GN/PN rates: 1.14 – 1.27) (**Figs. 4a,b and Supplementary Tables 7,8**). This effect is mediated by PN and GN mutation rates increasing and decreasing, respectively, as transcription levels increase (**Extended Data Fig. 5c**). Since regions of high gene expression (Q2, Q3, Q4) account for only 26% of the genome and 30% of interrogated base pairs on average while intergenic regions account for 62% of the genome and 57% of interrogated base pairs on average (**Extended Data Figs. 5a,b**), the relatively larger increase in PN mutation rates in regions of high expression is balanced by the relatively smaller increase in GN mutation rate in intergenic regions to yield similar total substitution mutation rates for PNs and GNs. Overall, these results suggest significant differences between PNs and GNs in the substitution mutagenesis effects of transcriptional activity and likely in their transcription-coupled repair.

**Figure 4.**
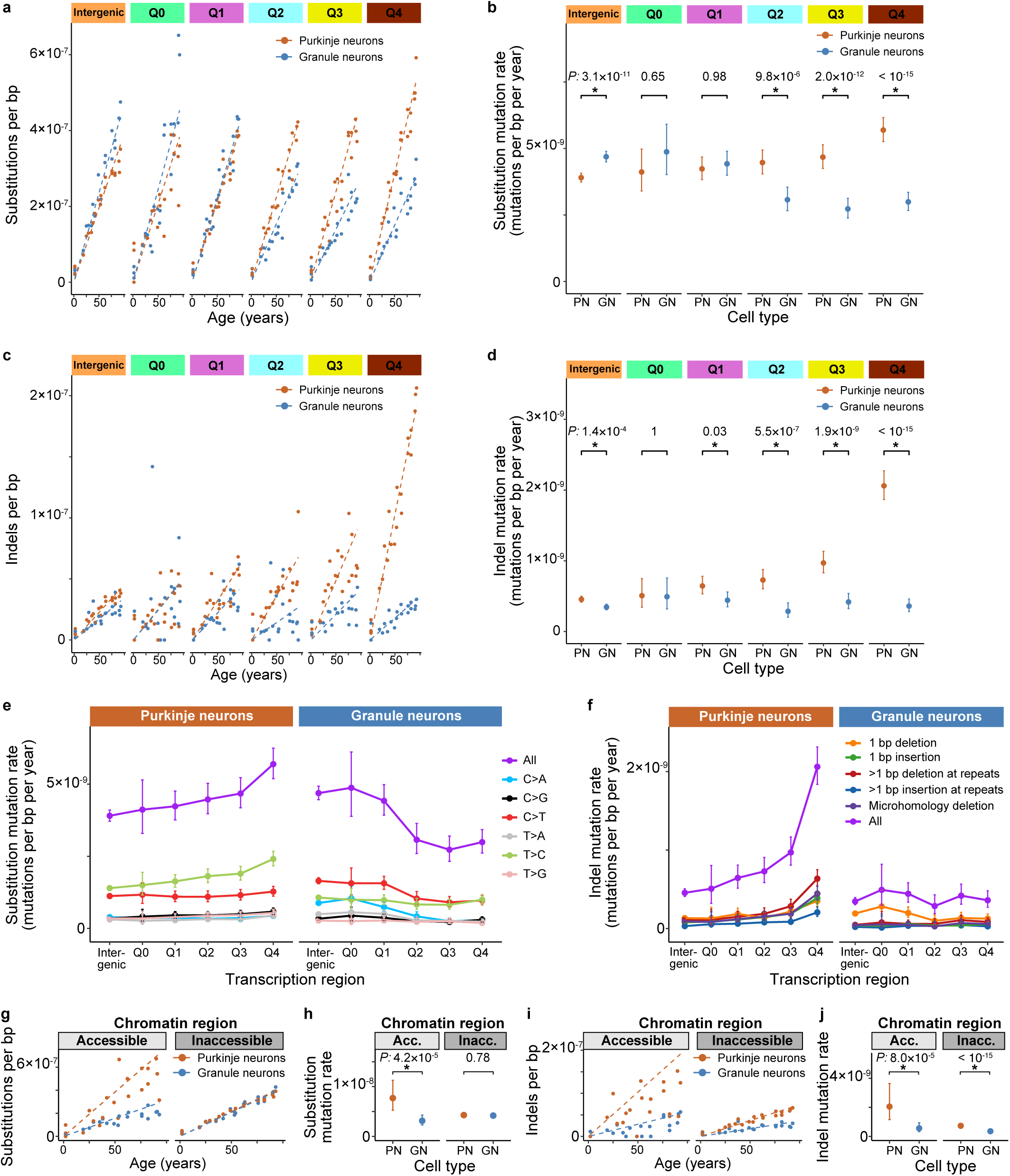
Associations of mutational processes with transcriptional activity and chromatin accessibility. **a-d,** Substitution (a) and indel (c) mutation burdens versus age, and comparisons between PN and GN substitution (b) and indel (d) mutation rates, for different transcription regions. Q0 is regions of genes that are not expressed, and Q1-Q4 are regions of expressed genes in bins of increasing expression quartiles. **e,f,** Substitution (e) and indel (f) mutation rates for different transcription regions, stratified by mutation type. Error bars show 95% CIs. See **Extended Data Figs. 5e,f** for plot of rate comparisons between PN and GN for each stratum. **g-j,** Substitution (g) and indel (i) mutation burdens versus age, and comparisons between PN and GN substitution (h) and indel (j) mutation rates (mutations/[bp·yr]), for different chromatin accessibility regions. Acc., accessible; Inacc., inaccessible. **a,c,g,I,** Dashed lines show predictions of the regression model. **b,d,h,j,** PN versus GN comparisons show *P*-values (asterisks mark significant differences) and error bars show 95% CIs. **a-j,** See **Supplementary Tables 7,8** for further details of rates and comparison statistics.

In contrast to the effects of transcription on substitutions, indel mutation rates were higher in PNs than in GNs in both intergenic regions and in every quartile of gene expression, with an especially large rate difference in the highest quartile (95% CI of PN/GN rates: Intergenic = 1.16 – 1.50, Q1 = 1.12 – 1.90, Q2 = 1.81 – 3.58, Q3 = 1.78 – 3.01, Q4 = 4.54 – 7.19) (**Figs. 4c,d and Supplementary Tables 7,8**). Moreover, in GNs, indel mutation rates did not vary significantly across intergenic, non-expressed genes, and expressed gene regions (**Extended Data Fig. 5d**). The observation that the mutation rate ratios of PNs versus GNs in transcribed regions is greater for indels than for substitutions (**Figs. 4b,d and Supplementary Table 8**) suggests that, in PNs, transcription has a relatively greater effect on indel mutagenesis than on substitution mutagenesis. Though, due to the higher overall mutation rate of substitutions compared to indels, transcription still contributes a larger absolute difference in mutation rate for substitutions than for indels (**Supplementary Table 7**). Together, these results indicate that the larger total indel mutation rate in PNs compared to GNs is primarily mediated by transcription, with an additional component due to an increased mutation rate in intergenic regions, and also that transcription has a minimal effect on indel mutagenesis in GNs.

We further characterized the effect of transcription by additionally stratifying by mutation type. This analysis showed that T>C mutations were the most notable substitution type increasing with transcription in PNs, while C>A, C>T, and T>A mutations were the most notable substitution types decreasing with transcription in GNs (**Fig. 4e and Supplementary Tables 7,8**). Moreover, all major indel types increased with transcription in PNs, whereas in GNs, the only notable trend was a decrease in the rate of 1 bp deletions with increasing transcription (**Fig. 4f and Supplementary Tables 7,8**). Additional comparisons of PN versus GN substitution and indel mutation rates across all transcription region x mutation type strata identified significant differences between PN and GN rates for nearly all mutation types in intergenic regions and in the highest quartile of gene expression (Q4), indicating that mutation patterns are most divergent between PNs and GNs in these regions (**Extended Data Figs. 5e,f and Supplementary Tables 7,8**). In contrast, almost none of the mutation types in non-expressed genic regions and only a subset of mutation types in lower quartiles of gene expression had divergent PN versus GN mutation rates (**Extended Data Figs. 5e,f and Supplementary Tables 7,8**). Further mutational signature analyses stratified by transcription region showed that SBS16-related burdens increased with gene expression level in both cell types and that ID4-, and ID11-related burdens were the most notable indel signatures increasing with gene expression level in PNs, consistent with these signatures being related to transcription^49,51^ (**Extended Data Figs. 6a,b and Supplementary Table 10**). ID5 and ID8 also increased mildly with transcription, suggesting they may also be affected by transcription in PNs (**Extended Data Figs. 6a,b and Supplementary Table 10**).

Next, we assessed for imbalances in mutation rates of each substitution mutation type between the transcribed versus the untranscribed strand (i.e., considering for each strand only those substitution mutations whose genome reference base at the site of the mutation is a pyrimidine). In both PNs and GNs, T>C / A>G mutations had the greatest strand asymmetry: T>C mutations occur in PNs and GNs at 1.3-fold and 1.7-fold higher rates, respectively, on the transcribed versus the untranscribed strand (**Extended Data Fig. 6c**). C>G / G>C mutations also exhibited concordant strand asymmetry in PNs and GNs, with 1.4-fold and 1.5-fold higher rates, respectively, in the untranscribed strand (**Extended Data Fig. 6c**). Notably, no mutation types exhibited discordant strand asymmetry between PNs and GNs (i.e., transcribed > untranscribed strand rate in one cell type with untranscribed > transcribed strand rate in the other cell type) (**Extended Data Fig. 6c**). The strand-stratified mutation spectra plotted across all trinucleotide contexts showed that the T>C mutation strand asymmetry is concentrated in ATN>ACN contexts (**Extended Data Fig. 6d**), consistent with the known contexts and transcriptional asymmetry of SBS16^52^ and similar to the pattern observed previously for cerebral cortex neurons^2^. Together, these results are consistent with our finding that T>C mutations exhibit the greatest difference in mutation rate between PNs and GNs (**Fig. 2d**), that T>C mutations have the greatest susceptibility to transcription level in PNs relative to other mutation types (**Fig. 4e**), and that there is a greater exposure to SBS16 in PNs than in GNs (**Fig. 3b**). Thus, the most divergent substitution mutation pattern between PNs and GNs is primarily mediated by transcription and concentrated in SBS16’s ATN>ACN contexts.

Transcription is closely coupled with chromatin state, and regions of accessible chromatin may be both more exposed to damage and more accessible to DNA repair^53,54^. We therefore compared mutation rates in accessible versus inaccessible chromatin regions based on prior single-cell chromatin accessibility data of human PNs and GNs^55^ (chromatin accessible regions: 0.7% and 3% of the genome and of interrogated base pairs on average in PNs and GNs, respectively) (**Extended Data Figs. 7a,b, Supplementary Table 11, and Methods**). This analysis showed that PNs have higher substitution mutation rates than GNs in accessible chromatin regions, while substitution mutation rates did not differ in inaccessible chromatin regions (95% CI of PN/GN rates: accessible chromatin = 1.62 – 3.71, inaccessible chromatin = 0.95 – 1.10) (**Figs. 4g,h and Supplementary Tables 7,8**). In contrast, PNs had higher indel mutation rates in both accessible and inaccessible chromatin regions (95% CI of PN/GN rates: accessible chromatin = 1.97 – 6.76, inaccessible chromatin = 1.78 – 2.34) (**Figs. 4i,j and Supplementary Tables 7,8**). Comparing rates between chromatin regions within each cell type showed both higher substitution and indel mutation rates in chromatin accessible than inaccessible regions in PNs (95% CI of accessible/inaccessible rates: substitutions = 1.28 – 2.53, indels = 1.67 – 4.68), but no rate differences in GNs (**Extended Data Figs. 7c,d and Supplementary Table 8**). Together, these results indicate that, similarly to transcription, chromatin accessibility is associated with higher substitution and indel mutation rates in PNs. Moreover, the total substitution mutation rates of PNs and GNs are similar despite PNs’ higher substitution mutation rates in accessible regions, because these regions account for only a small fraction of the genome. In contrast, an important contributor to the higher total indel mutation rate of PNs compared to GNs is the higher indel mutation rate of PNs not only in accessible, but also in inaccessible chromatin regions that comprise most of the genome.

## Discussion

Somatic mutation processes vary significantly across cell types, yet, to date, aging-related somatic mutation rates and patterns have only been profiled genome-wide in a very small fraction of the body’s cell types^2,46,56^. Identifying the factors driving cell type variability in somatic mutation processes may be essential for discovering the still enigmatic mechanisms by which most somatic mutations arise, as these factors are likely synonymous with or closely linked to the underlying causes of mutation. The gap in profiling somatic mutations at the level of cell types is especially significant for the brain that is comprised of a large diversity of cell types of which only a few have had genome-wide somatic mutation profiling^6,46^. Additionally, the remarkable diversity of cell types in the brain provides a rich landscape of physiological variability from which the key determinants and mechanisms of aging-related somatic mutation rates and patterns could be discovered. To the extent that genomic integrity plays a role in the vulnerability of specific brain cell types to specific neuropsychiatric diseases, as was established recently for Huntington’s disease^7^, characterizing somatic mutations at the level of cell types may also provide new strategies to intervene in these diseases.

Here, we developed a method to obtain cell type-specific somatic mutation profiles of PNs and GNs—the two major types of cerebellar neurons—that we chose due to their dramatically different characteristics and physiologies. This provided the first genome-wide landscapes of somatic mutation in the cerebellum, and we identified notable differences and surprising similarities in the somatic mutation processes of these two types of neurons that constrain hypotheses about drivers of aging-related somatic mutation.

Our most surprising finding was that the rate of substitution mutations was not significantly different between PNs and GNs, and only slightly different from that of cerebral cortex neurons in a prior study^2^. Our analyses of PNs and GNs showed that this was mainly due to the lack of a difference in the rate of the SBS5 mutational process to which most mutations were ascribed. This suggests that SBS5, which gives rise to most mutations throughout the body^56^, either arises via a mechanism that is largely invariant with cell size and metabolic rate, or that there exist homeostatic DNA repair or other mechanisms that are finely tuned to counteract deviations in the level of the SBS5 process. Another surprising finding was the low non-zero rate of SBS1 in both PNs and GNs, which was also recently observed in cerebral cortex neurons^46^. Recent studies have attributed SBS1 to cell division^47^, so this discrepancy may either be due to imperfect estimation of the low level of SBS1 exposures in our study and/or other studies, or that SBS1 is indeed active at low, yet non-zero levels in post-mitotic cells. Low-level contamination of our PNs and GNs with dividing cells is an unlikely explanation, as we observed this finding across two distinct and well-validated protocols that achieve high purity levels. Moreover, any putative cross-contamination would likely be random across samples such that SBS1 levels would not correlate with age.

Our data point to transcription as a key driver of the relatively small differences in substitution rates and patterns between PNs and GNs. Specifically, SBS16 levels, which are higher in PNs, increase with transcription as has been observed for cerebral cortex neurons^46^, and surprisingly, substitution rates increase in PNs but decrease in GNs at higher quartiles of gene expression. This latter finding parallels a prior study that found that cerebral cortex neurons have increased substitution rates while oligodendrocytes have decreased substitution rates at higher transcription quantiles^46^. We speculate that whether a cell type exhibits a relative increase or decrease of its substitution rate at its highest transcription quantiles depends on whether its highest transcription levels create transcription-coupled damage that exceeds (e.g., in PNs and cerebral cortex neurons) or is below (e.g., in GNs and oligodendrocytes) its capacity of transcription-coupled repair^54,57^.

Indels exhibited more significant relative differences in rates and patterns between PNs and GNs, which our analyses suggest is also primarily driven by transcription. The rates of all indel types increased with transcription levels in PNs, mutation rates were higher in PNs than GNs for all indels types except 1 bp deletions, and two of the indel signatures whose burdens increased with transcription levels were ID4 and ID11 that have been previously linked to transcription^49,51^. The higher burden of the ID8 signature in PNs may also be related to transcription, as ID8 levels increased with transcription level and ID8’s association with DNA double-strand breaks^44^ is consistent with prior studies that identified transcription-induced double-strand breaks in neural cells^58^. Interestingly, in ataxia telangiectasia, a genetic disorder characterized by deficient double-strand break repair, PNs are more susceptible to degeneration^59^. PNs also had higher indel rates in intergenic regions, which may be due to background transcription of intergenic regions^60^, or due to non-transcription-related process.

Our study has a few limitations. First, our duplex sequencing used short sequencing reads that preclude profiling mutational patterns at sites of long repeats, such as those in which pathogenic mutations cause neurodegeneration. Those sites may have distinct mutational processes than those we have profiled^7^. Second, de novo signature extraction is challenging with even a few dozen samples^61^ so that our estimates of signature exposure relied on known COSMIC signatures. As duplex-sequencing methods emerge that dramatically increase the yield of somatic mutations^62^, signature extraction in smaller sample sets will become more reliable, though scaling to larger numbers of samples will still be important. Third, our process for purifying PNs with LCM could feasibly be combined with immunostaining to profile many other cell types, but it is laborious. A notable recent preprint developed duplex sequencing of accessible chromatin for droplet-encapsulated single cells along with concomitant RNA-sequencing to profile somatic mutations in several neuronal and non-neuronal cell types in one experiment^63^, pointing at a way forward to scale cell type-specific mutation profiling. However, more rare cell types may still require other methods of purification.

Overall, our data support the importance of conducting future systematic cell type-resolved somatic mutation studies. Delineating somatic mutations at the level of cell types in disease states, for example in neurodegenerative diseases, will also be important for understanding cell type-specific vulnerabilities in maintaining genomic integrity under stress. As somatic mutation landscapes are defined for many more cell types, combining these data with other cell type-resolved multi-omic data will shed new light on the origins of somatic mutations.

## Methods

### Samples

Human post-mortem cerebellum, cerebral cortex, liver, and kidney of individuals with no known neurological or psychiatric disease were obtained from the NIH NeuroBioBank (University of Maryland, Mount Sinai, and the Human Brain and Spinal Fluid Resource Center sites). Since cerebellar development continues postnatally, we selected brains from individuals at least 2 years old^64^. For cerebellum, we sampled from the lateral cerebellar hemisphere. For cerebral cortex, we sampled from the temporal lobe (Brodmann area 20). Individuals whose cerebellar tissue after staining was found to have low quality (i.e., significant freezing artifacts and/or morphological distortion) were excluded from the study prior to performing laser-capture microdissection (LCM). See **Supplementary Table 1** for details of individuals included in the study.

### DNA extraction from germline reference tissues

DNA was extracted from approximately 25 mg of tissue (see **Supplementary Table 1** for a list of which tissue was used for the germline reference for each individual) using the MagAttract HMW DNA Kit (Qiagen) per the manufacturer’s protocol for tissues. DNA was eluted with 150 µL of 10 mM Tris, pH 8.0 (dilution made from Invitrogen, AM9855G). The concentration and quality of DNA samples was measured using a NanoDrop instrument (Thermo Fisher Scientific), a Qubit 1× dsDNA HS Assay Kit (Thermo Fisher Scientific), and a Genomic DNA ScreenTape TapeStation Assay (Agilent). DNA was then stored at −20 °C.

### Cerebellum cryosectioning and staining

The cryosectioning and staining procedure was adapted from a prior protocol^65^. Before cryosectioning, the cryostat (Leica CM1950) was loaded with a new blade, and the cryostat working area was cleaned with RNaseZap (Thermo Fisher, AM9780) and ethanol. An approximately 10 mm x 10 mm x 5 mm piece of cerebellum was mounted with Tissue-Tek O.C.T. compound (Sakura, 4583) onto the cryostat specimen disc and left in the cryostat until hardened. Then 10 µm tick sections were cut onto glass PEN-membrane slides (Leica, 11505189), dried for 10 min at room temperature, and stained with hematoxylin and eosin as follows (each step took place in a separate container): (1) 2 min in 70% ethanol (Decon Labs, 8601), then wiping off excess ethanol; (2) 10 dips (each dip is ∼ 1 sec) in water from a Picopure system (Hydro Services), then repeating this a second time in a second water container; (3) 5 min in hematoxylin (Epredia, 7211), then wiping off excess hematoxylin; (4) 1 min in Picopure water with dipping every few seconds, then repeating this two more times each in a new water container; (5) 2 dips (each dip is ∼ 1 sec) in 70% ethanol; (6) 3 dips in eosin (Leica, 3801619), then wiping off excess eosin; (7) 3 dips in 70% ethanol; (8) 10 sec in 70% ethanol; (9) 10 sec in 100% ethanol (Decon labs, 2701), then repeating this a second time in a second 100% ethanol container; (10) air drying until no ethanol residue is visible. The slides were then either taken directly to LCM or placed in a slide box within an airtight plastic bag and stored at −80 °C.

### Laser-capture microdissection of Purkinje neurons for DNA sequencing

Before use, LCM equipment was cleaned with DNAzap (Invitrogen, AM9890). LCM samples were cut with a Leica LMD6000 instrument equipped with Leica Laser Microdissection software V8.0.1 and a 40X objective. For each sample, the laser was calibrated using the 40X objective, and we set the laser power to 16, aperture to 2, and cutting speed to 7. Purkinje cell bodies were selected manually using the 40X objective and cut in “Draw+Cut” mode. For each individual, approximately 600 – 1200 PN cell bodies (see **Supplementary Table 1** for numbers for each sample) were cut into the cap of a Low Binding 0.65 mL tube (Costar, 3206) containing 55.5 µL of 30 mM Tris-HCl, pH 8.0 and then centrifuged at 21,130 x g at 4 °C for 5 min. We then added to each sample 1.5 µL of 20% Tween-20 (dilution made from Promega, H5152), 1.5 µL of 20% IGEPAL CA-630 (dilution made from Sigma, I8896), and 1.5 µL of Thermolabile Proteinase K (NEB, P8111S). The volume of each tube was then adjusted to a total of 60 µL with nuclease-free water (NFW; Thermo Fisher, AM9938) by comparing the volume to a control tube containing 60 µL NFW to compensate for evaporation that occurred during LCM, and then briefly spun down with a counter-top mini-centrifuge. Samples were then mixed at 1,000 rpm for 1 min at room temperature on a ThermoMixer C (Eppendorf), centrifuged at 21,130 x g at 4 °C for 1 min, and incubated at 37 °C and 1,000 rpm for 1 hour in a ThermoMixer C. Samples were then briefly spun down with a counter-top mini-centrifuge, transferred to 0.2 mL low-binding tubes (Axygen, PCR-02-L-C), and incubated in a thermocycler at 55 °C for 10 min to deactivate the Proteinase K. The samples were then either frozen at −20 °C or taken directly to NanoSeq library preparation.

### Laser-capture microdissection for RNA-sequencing

Additional LCM was performed as described in the above LCM for DNA sequencing protocol for three individuals (IDs: 937, 5609, and 11589) whose tissues had RNA integrity values ≥8.0 per the NIH NeuroBioBank, except: (1) the LCM instrument was cleaned with RNaseZap instead of DNAzap; (2) Approximately 300 PN cell bodies per sample were cut, and separately approximately 250 - 300 circles of ≍ 20 µM diameter each from the granule cell layer (each circle containing a cluster of about 10 granule neurons) per sample were cut, into the cap of a 0.65 mL Low Binding tube containing 60 µL of lysis buffer containing: 10 µL of 40% Poly-ethylene Glycol 8000 (Sigma, 89510-250G-F), 0.6 µL of 10% Triton X-100 (dilution made from Promega, H5142), 0.75 µL of 40 U/µL RNase Inhibitor (NEB, M0307), 0.4 µL of 100 µM OligodT30VN (/5Biosg/ACGAGCATCAGCAGCATACGATTTTTTTTTTTTTTTTTTTTTTTTTTTTTTVN, where ‘N’ is a mixture of all 4 bases and ‘V’ is a mixture of A, C, or G; IDT, ordered with HPLC purification), 4 µL of 10 mM dNTP mix (NEB, N0447S), and 44.25 µL NFW. After LCM, samples were centrifuged at 21,130 x g at 4 °C for 5 min, and the volume of each tube was adjusted to 60 µL NFW. Samples were centrifuged at 21,130 x g at 4 °C for 1 min, frozen in −80 °C for 15 min to further lyse cells, and thawed on ice for 5 min. Samples were then incubated at 72 °C for 10 min, immediately placed on ice, and then centrifuged at 21,130 x g at 4 °C for 1 min. Sample were then immediately taken to RNA library preparation.

### Purification of granule neuron nuclei by fluorescence-activated nuclei sorting for DNA sequencing

Approximately 20 mg of cerebellum was cut and added to 4.5 ml of chilled lysis buffer (0.32 M sucrose, 5 mM CaCl_2_, 3 mM magnesium acetate, 0.1 mM EDTA, 1 mM DTT, 10 mM Tris HCl pH 7.5, 0.1% Triton X-100) in a dounce homogenizer (DWK, 885300-0007). The tissue was dounced on ice 25 times with pestle size A and then 25 times with pestle size B. The homogenate was passed through a 30 µm MACS SmartStrainer filter (Miltenyi Biotec, 130-098-458) and then layered on a 7.4 mL sucrose cushion (1.8 M sucrose, 3 mM magnesium acetate, 1 mM DTT, 10 mM Tris HCl pH 7.5) in a chilled ultracentrifuge tube (Seton Scientific, 7030). The tubes were centrifuged at 10,000 rpm (maximum 17,000 x g) at 4 °C for 1 h using a TH-641 rotor in a Sorvall wX+ ultra-centrifuge (Thermo Fisher Scientific). The supernatant was discarded, and 1 mL of chilled nuclei resuspension buffer (0.25 M sucrose, 5 mM calcium chloride, 3 mM magnesium acetate, 0.1 mM EDTA, 10 mM Tris-HCl pH 7.5) was placed on top of the nuclei pellet for 10 min while on ice. The pellet was then resuspended with wide-bore 1000 µL tips and transferred to a 5 mL tube. An additional 1 ml of nuclei resuspension buffer was added to the sample and the nuclei were further mixed 20 times with wide-bore 1000 µL tips. The nuclei were again passed through a 30 µm MACS SmartStrainer filter. Nuclei were imaged on a hemocytometer with a light microscope after staining with a 0.2% final concentration of trypan blue (Thermo Scientific, 15250061), and for samples with many nuclei clumps, we performed additional pipette mixing 30 to 50 times with wide-bore 1000 µL tips and passed the nuclei again through a 30 µm MACS SmartStrainer filter.

We measured the volume of the nuclei suspension, we added an equal volume of 2% PFA fixation solution (0.25 M sucrose, 5 mM calcium chloride, 3 mM magnesium acetate, 0.1 mM EDTA, 10 mM Tris-HCl pH 7.5, 2% PFA [from Thermo Fisher, 28906]) to achieve a final concentration of 1% PFA, and we placed the sample on a rotator for 10 min at room temperature. We then added glycine quenching solution (2.5 M glycine [Sigma-Aldrich, 50046] in NFW) to the fixed nuclei to a final concentration of 0.2 M glycine, and incubated on a rotator at room temperature for 10 min. The nuclei were centrifuged at 1,000 x g at 4 °C for 5 min, the supernatant was discarded, the nuclei pellet was resuspended with 1 mL of staining buffer (2% bovine serum albumin [VWR, 0332-25G], 1X PBS [from 10X PBS, Thermo Scientific, AM9625], 3 mM MgCl_2_, 0.1% Tween-20 [Promega, H5152]), and the nuclei were transferred to a 1.5 mL tube and placed on a rotator at 4 °C for 30 min. Nuclei clumps were broken up by pipette mixing 100 times with wide-bore 1000 µL tips, followed by an additional 100 to 500 times with standard 1000 µL tips. For samples with residual clumps, we filtered the nuclei again once or twice using a 30 µm MACS SmartStrainer filter.

We then added 1.2 µg of Alexa Fluor 647 anti-NeuN antibody (Abcam, ab190565) to each sample and incubated it on a rotator at 4 °C for 1 h. Nuclei were centrifuged at 1,000 x g at 4 °C for 5 min, the supernatant was discarded, and the nuclei were resuspend with 500 µL wash buffer (1X PBS, 3 mM MgCl_2_, 0.1% Tween-20) using wide-bore 1000 µL tips followed by 30 to 100 times with 200 µL tips, and then incubated on a rotator at 4 °C for 5 min. Nuclei were centrifuged again at 1,000 x g at 4 °C for 5 min, the supernatant was discarded, and the nuclei were resuspended with 1000 µL FACS buffer (1X PBS, 3 mM MgCl_2_) using wide-bore 1000 µL tips. For samples with residual clumps based on microscopy imaging, we pipette mixed 200 to 600 times with standard 1000 µL tips and/or passed the sample through a MACS SmartStrainer filter. For all samples, we then passed the sample through a 30 µm MACS Pre-Separation filter (Miltenyi Biotec, 130-041-407).

Granule neuron nuclei (18,000 nuclei per sample) were sorted using a SONY SH800 instrument using the ‘NeuN(+) low FSC, low BSC, singlet gate’ shown in **Extended Data Fig. 1b** into 42 µL of sort buffer (857 mM Tris pH 8.0, 57 mM NaCl) in a 1.5 mL DNA LoBind tubes (Eppendorf). We did not sort ‘NeuN(+) high FSC’ events, as these are multiplets of granule neurons (**Extended Data Figs. 1b,c**). Note, the sort buffer composition was chosen so that during the subsequent formalin cross-link reversal reaction in a volume of 120 µL this would yield a final concentration of 300 mM Tris pH 8.0, 20 mM NaCl. After sorting, sample tubes were centrifuged at 500 x g at 4 °C for 5 min and frozen at −20 °C.

### Fluorescence-activated nuclei sorting for single-nucleus RNA-sequencing

Nuclei were prepared from two cerebellum of two individuals (937 and 11589) as described in the above section ‘Purification of granule neuron nuclei by fluorescence-activated nuclei sorting for DNA sequencing’, with the following changes: (1) ∼100 mg of tissue was used; (2) the lysis buffer contained 0.4 U/µL RNase Inhibitor (Sigma, 3335402001) and it did not contain Triton X- 100; (3) before the final MACS SmartStrainer filtering prior to fixation, we added an additional 400 µL of nuclei resuspension buffer; (4) the nuclei were fixed by adding an equal volume of 8% PFA fixation solution and incubating for 1 h; (5) nuclei were stained with 2.2 µg of anti-NeuN antibody; (6) the final filtering before sorting was performed with a MACS SmartStrainer filter; (7) a maximum number of nuclei were sorted using the ‘NeuN(+) low FSC, low BSC, singlet gate’ and the ‘NeuN(-)’ gate shown in **Extended Data Fig. 1b** into a 5 mL FACS tube (Corning, 352063) containing 1 mL of sort buffer (5% BSA, 1X PBS, 3 mM MgCl_2_); note, the FACS tube was coated prior to sorting with 5 mL of sort buffer for 10 min.

### NanoSeq library preparation and sequencing

The below protocol was based on the previously published NanoSeq protocol^2^, which we optimized for lower DNA input via two key changes: (1) using a ligase optimized for low DNA input, and, (2) when possible, keeping the DNA library with the SPRI (solid-phase reversible immobilization) beads after the elution step of SPRI bead purifications (i.e., ‘on-bead’ library preparation).

#### SPRI beads and PEG solution preparation

SPRI beads used in the library preparation were made by washing 1 ml Sera-Mag carboxylate-modified SpeedBead (Cytiva, 65152105050250) and resuspending the beads in 50 ml of 18% PEG-8000, 1.75 M NaCl, 10 mM Tris pH 8, 1 mM EDTA, and 0.044% Tween-20. PEG solution used in the library preparation was obtained by placing the prepared SPRI beads on a magnet and transferring the PEG solution without the beads to a new tube.

#### Sample preparation

For germline reference samples, 50 ng of DNA was first diluted in a 0.2 mL low-binding tube with NFW to a total volume of 60 µl.

For germline reference and LCM PN samples, DNA was then purified by adding SPRI beads at a 2.3X SPRI bead to sample volume ratio, followed by two 80% ethanol washes. DNA was eluted from the beads with 20 µL NFW, and the eluted DNA was kept with the beads rather than transferring to a new tube.

For sorted GN samples, formaldehyde cross-linking reversal and DNA extraction was first performed by adding 12 µL of Proteinase K (Qiagen, 19131) and 12 µL of 10% SDS (Thermo Scientific, AM9822) to the ∼ 96 µL volume of sorted nuclei, vortexing for 20 sec, and incubating at 56 °C overnight (∼15 h). We then added 112 µL of Buffer AL (Qiagen, 19075) and 8 µL of Proteinase K (Qiagen, 19131), vortexed for 20 sec, and incubated at 56 °C for 1 hour. Samples were placed on ice for 3 min and then at room temperature for 5 min. DNA was then purified by adding diluted SPRI beads (SPRI beads diluted with an equal volume of PEG solution) at a 1.5X diluted SPRI bead to sample volume ratio, followed by two 80% ethanol washes. DNA was eluted with 25 µL of NFW, and 20 µl of the elution was transferred to a new tube.

#### Library preparation

For all sample types, DNA was fragmented by adding a 5 µl fragmentation mix comprised of 2.5 µl of either 10X CutSmart Buffer (NEB, B7204S) or 10X rCutSmart Buffer (NEB, B6004), 2 µl NFW, and 0.5 µl of HpyCH4V (NEB, R0620S). Reactions were incubated at 37 °C for 15 min.

Reactions were cleaned up by adding a 2.5X PEG solution to sample volume ratio (germline reference and LCM PN samples) or 2.5X SPRI beads to sample volume ratio (sorted GN samples), with two 80% ethanol washes, and eluting the DNA with 20 µL of NFW, keeping the eluted DNA with the beads.

The 20 µL of fragmented DNA was A-tailed by adding a 10 ul A-tailing mix comprised of 3 µl of 10X NEBuffer 4 (NEB, B7004S), 3.7 µl NFW, 3 µl of 1 mM each dATP/ddCTP/ddGTP/ddTTP (dATP/ddBTP) (dATP, Thermo Fisher Scientific, R0141; ddCTP/ddGTP/ddTTP, Jena Bioscience NU-1019S), and 0.3 µL of Klenow fragment 3′→5′ exo-(NEB, M0212S, 5U/µL). Reactions were incubated at 37 °C for 30 min.

The 30 µL of A-tailed DNA was ligated to adapters by adding to each sample 15 µl 4X Ultralow Input Ligation Buffer 4X (Qiagen, 180492), 3 µl of Ultralow Input Ligase (Qiagen, 180492), and 0.88 µL (germline reference samples) or 0.48 µL (LCM PN and sorted GN samples) of xGen CS Adapter (IDT, 1080799, 15 µM stock concentration), and NFW to a total volume of 60 µL. Reactions were incubated at 25 °C for 30 min. Reactions were cleaned up by adding a 1X PEG solution to sample volume ratio (germline reference and LCM PN samples) or 0.9X PEG solution to sample volume ratio (sorted GN samples), with two 80% ethanol washes, and the DNA was eluted with 46 µL of NFW, keeping the eluted DNA with the beads. A second clean-up was performed to remove residual adapter dimers by adding a 1X PEG solution to sample volume ratio (germline reference and LCM PN samples) or 0.8X PEG solution to sample volume ratio (sorted GN samples), with two 80% ethanol washes, and the DNA was eluted with 20 µL of NFW (germline reference and LCM PN samples) or with 46 µL of NFW (sorted GN samples), keeping the eluted DNA with the beads.

Ligated DNA was quantified for each sample by quantitative PCR (qPCR) as previously described^2^ using a QuantStudio 6 Flex instrument (Applied Biosciences) and using as input 4 µL of 1:5,000 dilutions for germline reference samples, 1:500 dilutions for LCM PN samples, and 1:1,000 dilutions for sorted GN samples (dilutions were made in 10 mM Tris-HCl pH 8.0, 0.05% Tween-20).

We then performed PCR amplification of libraries with input DNA amounts based on qPCR quantification. We performed PCR amplification of LCM PN samples once for initial sequencing, and then a second time to obtain additional deeper sequencing. We input into the PCR reactions for germline reference samples 12 fmol of DNA, for LCM PN samples either 0.6 fmol (initial libraries) or 1.2 fmol (additional libraries for deeper sequencing; except for individuals 1791, 1819, and 2794 for whom we input the maximum quantity we had available for deeper sequencing: 1 fmol, 0.6 fmol, and 0.2 fmol, respectively), and for sorted GN samples 1.8 fmol. PCR reactions were comprised of the input DNA, 25 µL NEBNext Ultra II Q5 Master Mix (NEB, M0544S), 5 µL of xGen UDI primers (IDT, 10005922), and NFW to a total volume of 50 µL. Reactions were incubated at 98 °C for 30 sec; [98 °C for 10 sec; 65 °C for 75 sec] X 9 cycles for 12 fmol inputs / 12 cycles for 0.6 fmol inputs / 11 cycles for additional LCM PN libraries performed for deeper sequencing and for sorted GN libraries; 65 °C for 5 min. Reactions were cleaned up by adding a 0.9X SPRI beads solution to sample volume ratio (germline reference and LCM PN samples) or 0.7X SPRI beads solution to sample volume ratio (sorted GN samples), with two 80% ethanol washes, and the DNA was eluted with 22 µL of NFW (germline reference and LCM PN samples) or with 48 µL NFW (sorted GN samples), transferring the eluted DNA to a new tube. Sorted GN samples were cleaned up a second time by adding a 0.7X SPRI beads solution to sample volume ratio, with two 80% ethanol washes, and the DNA was eluted with 24 µL of NFW, transferring the eluted DNA to a new tube. Final libraries were evaluated using a High Sensitivity D5000 ScreenTape TapeStation Assay (Agilent).

#### Sequencing

Libraries were sequenced on NovaSeq 6000 (some of the data for PNs from individuals 5559, 5344, and 4643) and NovaSeq X (all remaining data) instruments (Illumina) with 150 bp paired-end reads, targeting a depth of 90 Gb per 0.6 fmol DNA input into the PCR reaction. See **Supplementary Table 2** for sequencing metrics.

### RNA library preparation and sequencing of laser-capture microdissection samples

Smart-seq3 was performed as previously described^66^ with the following modifications: 1) the reverse transcriptase reaction used Tris-HCl pH 8.4 (Teknova, T1084); 2) the reverse transcriptase reaction volume was scaled to a total volume of 80 µL to accommodate the input volume from LCM; 3) preamplification PCR was performed in a volume of 110 µL with 20 cycles; 4) samples were frozen at −20 °C after cDNA purification, and processing continued the following day; 5) 2 ng of cDNA was used for the tagmentation reaction; 6) the tagmentation PCR mix also contained 0.01% Tween-20, and the tagmentation PCR reaction was performed in a total volume of 20 µL with 15 cycles. Libraries were sequenced on a NovaSeq 6000 and NovaSeq X instruments with 150 bp paired-end reads. See **Supplementary Table 2** for sequencing metrics.

### Single-nucleus RNA library preparation and sequencing of nuclei purified by fluorescence-activated nuclei sorting

Single-nucleus RNA libraries were prepared using the GEM-X Flex Gene Expression Human kit (10x Genomics, 1000792) per the manufacturer’s instructions with the following parameters: (1) 300,000 nuclei of ‘NeuN(+), low FSC, low BSC, singlet’ gate samples and 200,000 nuclei of ‘NeuN(-)’ gate samples were input into the hybridization reaction; (2) the probe barcode was BC008; (3) the target recovery was 15,000 nuclei per sample; (4) to remove a peak at ∼ 190 bp of single unligated probe that we observed after an initial final library PCR, we added an additional cleanup of pre-amplification libraries by adding 30 µL of NFW to 40 µL of bead-cleaned pre-amplification library, adding 1.8X SPRIselect beads to sample volume ratio, with two 80% ethanol washes, eluting with 21 µL of NFW, and using 20 µL of eluted library for the indexing PCR; (5) the indexing PCR was performed with 14 cycles.

Libraries were sequenced on a NovaSeq X instrument with 28 bp (read 1) x 90 bp (read 2) reads. See **Supplementary Table 2** for sequencing metrics.

### Smart-seq3 data analysis

We used zUMIs^67^ v2.9.7 (https://github.com/sdparekh/zUMIs) to analyze Smart-seq3 data. We first created a transcriptome reference using the STAR^68^ genomeGenerate command and the GENCODE v32 release GRCh38 primary assembly transcriptome annotation file (gencode.v32.primary_assembly.annotation.gtf). Next, we prepared the input files by concatenating the fastq files of each sample and then processing them with the zUMIs merge_demultiplexed_fastq.R script. Next, we ran zUMIs with a YAML configuration file containing the following parameters: (1) R1 fastq base_definition: cDNA(23-150); UMI(12-19); (2) R2 fastq base_definition: cDNA(1-150); (3) index fastq base_definition: BC(1-8); (4) reference STAR_index and GTF_file parameters corresponding to the above transcriptome reference; (5) additional_STAR_params: --clip3pAdapterSeq CTGTCTCTTATACACATCT; (6) filter_cutoffs: BC_filter num_bases = 3 and phred = 20; UMI_filter num_bases = 2 and phred = 20; (7) barcodes: automatic = no, BarcodeBinning = 0, nReadsperCell = 100, and demultiplex = yes; (9) counting_opts introns = yes, downsampling = ‘0’, strand = 0, Ham_Dist = 1, velocyto = no, primaryHit = yes, and twoPass = no; (10) make_stats: yes.

We obtained the per-gene count of UMIs that aligned to exons from the umicount$exon$all data in the dgecounts.rds file produced by zUMIs. For each sample, we normalized its UMI counts by dividing the per-gene UMI counts by the total UMI counts of the sample and multiplying by 1,000,000. The heatmap was plotted with the ComplexHeatmap package^69^.

### Single-nucleus RNA-sequencing data analysis

Raw sequencing reads were first demultiplexed and processed using Cell Ranger software v.9.0.1 (10x Genomics) with the ‘multi’ function with default settings. In R^70^, we used Seurat v.5.1.0 to analyze the data. We first loaded the data from the ‘count/sample_filtered_feature_bc_matrix’ folders created by Cell Ranger for each sample into a single Seurat object. We then filtered to keep nuclei with between 200 to 5000 detected transcribed genes, between 250 to 10,000 UMI read counts, and a maximum of 5% mitochondrial reads. See **Supplementary Table 2** for basic nuclei counts and other metrics.

Next, we used the ‘split’ function to divide the Seurat object by individual (i.e. 937 and 11589), and we applied the ‘SCTransform’ and ‘RunPCA’ functions to the Seurat object. We then integrated the data across the two individuals using the ‘IntegrateLayers’ function with the settings ‘method = CCAIntegration, normalization.method = “SCT”, orig.reduction = “pca”, new.reduction = “integrated.cca”, dims = 1:30’. Next, we applied to the Seurat object the following functions: (1) ‘FindNeighbors’ with the settings: ‘dims = 1:30, reduction = “integrated.cca”, k.param = 20’; (2) ‘FindClusters’ with the settings: ‘cluster.name = “integrated.cca.clusters”, resolution = 0.8’; (3) ‘RunUMAP’ with the settings: ‘reduction = “integrated.cca”, reduction.name = “umap.cca”, dims = 1:30, n.neighbors = 30, min.dist = 0.1, spread = 0.5’. We then sub-clustered clusters that spanned > 1 UMAP visual cluster using the ‘FindSubCluster’ function with the settings: ‘graph.name = “SCT_snn”, subcluster.name = “integrated.cca.clusters.subclustered”’ and ‘resolution’ set to between 1 and 4 depending on the cluster being analyzed. The resulting cluster IDs were reordered numerically by similarity using the ‘BuildClusterTree’ function with the settings ‘dims = 1:30, reduction = “integrated.cca”, reorder = TRUE, reorder.numeric = TRUE’. We then merged clusters that were co-located visually in the same UMAP cluster.

Differential gene expression analysis was performed by applying to the Seurat object the function ‘JoinLayers’ with the setting ‘assay=”RNA”’, the function ‘PrepSCTFindMarkers’, and the function ‘FindAllMarkers’ with the setting ‘assay = “SCT”’. We assigned cell types to clusters using as a reference the Human Brain Cell Atlas^71^ cerebellar vermis, deep nuclei, and lateral hemisphere datasets available at the CZI CellxGene website (https://cellxgene.cziscience.com), another prior snRNA-seq study of human cerebellum^72^, and prior published reviews^73,74^.

Clusters representing multiplets of two or more different cell types were identified by their co-expression of markers characteristic of more than one cell type. The final UMAP and gene expression dot plot were made using Seurat’s DimPlot and DotPlot functions and with ggplot2^75^.

### NanoSeq data analysis

#### Primary data processing

NanoSeq primary data processing was performed using the NanoSeq analysis pipeline v.3.5.5 (https://github.com/cancerit/NanoSeq)^2^ for chromosomes 1–22, X, and Y (hg38 reference genome). We used the following non-default settings— ‘cov’ step: --exclude chrM,%random,chrUn_%,%_alt,chrEBV,HLA%; ‘var’ step: -c 0, ‘indel’ step: -c 0 --t3 135 --t5 10. In the ‘dsa’ step, we used the SNP mask ‘SNP.sorted.GRCh38.bed.gz’ supplied by the creators of the NanoSeq analysis pipeline (link in the above GitHub page) and a NOISE mask that we created as described in ref. ^45^. Following primary data processing, we used VerifyBamID2^76^ to exclude significant contamination from an unrelated sample as described in the NanoSeq GitHub page. We also evaluated the computationally estimated error rates output by the NanoSeq pipeline in the total_error_rate field of the results.estimated_error_rates.tsv files. The estimated error rate for each sample, which is the probability of a false-positive single-base substitution per bp, is calculated by the NanoSeq pipeline using the observed frequencies of single-strand consensus calls; i.e., as the probability of two reverse complement single-strand consensus calls summed across all trinucleotide contexts. Across PN and GN samples, this maximum estimated error rate was 3.0 × 10^-10^ and 1.6 × 10^-11^, respectively, confirming that both LCM and formalin fixation used during the purification of the respective sample types do not contribute a significant burden of false positive mutation calls relative to the total mutation burdens (**Supplementary Table 2**).

#### Fingerprinting

To further confirm that all the NanoSeq samples of each individual corresponded to the same individual, we performed a fingerprinting analysis as described in ref. ^45^. Briefly, this analysis counted the number of high-quality germline variants detected in each individual’s germline reference sample that were absent from each NanoSeq somatic sample. Somatic samples that originate from the same individual as the germline reference sample would be expected to have very few or none of the high-quality germline variant calls absent, assuming the variant locus had sufficient sequencing depth in the somatic sample.

#### Data analysis

We used our R package NanoSeqTools^45^ (https://github.com/evronylab/NanoSeqTools) to load all the NanoSeq pipeline output files required for downstream analysis. The load_nanoseq_data function of NanoSeqTools was run with bedtools_bin set to BEDTools v2.31.1^77^ and with exclude_regions set to the union of (1) the SNP and NOISE masks described above for NanoSeq primary data processing, and (2) an additional mask of regions we identified as having artifacts after analyzing the output of NanoSeq somatic mutation calls. This additional mask filtered only 33 bases in 18 loci, and it was created by identifying bases in which a somatic mutation was called in three or more samples in the study, and bases in which two or more individuals each had more than one somatic mutation call at that base.

Analysis of mutation burdens according to gene expression and transcribed strand utilized this study’s cerebellum snRNA-seq data as a reference for PN and GN gene expression. Specifically, we extracted from this data pseudo-bulk gene expression values for the PN and GN cell type clusters using the AggregateExpression function of Seurat. We used this data and the Ensembl v113 Homo sapiens gene annotation to then extract six sets of genomic regions for each cell type: intergenic regions, regions of genes that are not expressed (i.e., expression = 0; region set labeled Q0), and regions of genes that are expressed (i.e. expression > 0) binned in four quartiles of gene expression (region sets labeled Q1-Q4). We then removed from each of these region sets those regions that are annotated in the Ensembl annotation as being transcribed in both strands or that are present in more than one region set. Next, we used the NanoSeqTools package’s load_nanoseq_regions function to extract mutation data for each cell type using that cell type’s set of genomic regions. For substitution mutations in each of the five genic regions, we extracted mutations separately depending on whether the mutated pyrimidine (<u>C</u>>A, <u>C</u>>G, <u>C</u>>T, <u>T</u>>A, <u>T</u>>C, <u>T</u>>G) was on the transcribed or on the untranscribed strand. Overall, for each sample, this yielded 11 sets of substitution mutation data (intergenic mutations plus transcribed and untranscribed mutations for each of 5 genic regions) and 6 sets of indel mutation data (intergenic mutations plus each of 5 genic regions). The list of regions within each gene expression region set are listed in **Supplementary Table 11**.

Analysis of mutation burdens according to chromatin state utilized a prior study’s single-cell chromatin accessibility data for human cerebellum^55^. Specifically, we downloaded that study’s annotation of chromatin accessible regions (termed by the study as candidate *cis*-regulatory elements) from the CATlas website (https://decoder-genetics.wustl.edu/catlasv1/catlas_downloads/humanbrain/cCREs/cCREs.bed), filtering by cell-type = ‘PKJ_1’ or ‘PKJ_2’ to obtain PN data and by cell-type = ‘CBGRC’ to obtain GN data, and then padding these regions by 250 bp on each side. Chromatin inaccessible regions for each cell type were defined as genomic regions outside chromatin accessible regions. Finally, we used the NanoSeqTools load_nanoseq_regions function to obtain mutation data for each cell type for each of the two types of chromatin regions. The list of regions within each chromatin accessibility region set are listed in **Supplementary Table 11**.

To analyze mutation rates across different covariates such as cell type and mutation type, we fit a generalized linear model with the glmmTMB package^78^ with a negative binomial type 1 family to account for overdispersion that we observed with a Poisson model in residual plots and by the DHARMa package^79^. The model utilized a “log” link function, and the model formula was specified as: *# of mutations* ∼ *Main Covariate* ∗ *Secondary Covariate 1* (if specified) ∗ *Secondary Covariate 2* (if specified) ∗ … + offset(log( *Exposure* ))’. *Exposure* was calculated as *Age* × *# of Interrogated bp* × *# of mutations* / *# of mutations corrected for the trinucleotide distribution of interrogated base pairs relative to the genome*^ref.2^ when *# of mutations* = 0, and otherwise, as *Age* × *# of Interrogated bp*. The dispersion parameter was modeled as a function of mean-centered log(*Exposure*) by setting the parameter dispformula = ∼ I(log(*Exposure*) - mean(log(*Exposure*))) ), which allows overdispersion to vary with exposure. Main and secondary covariates varied by analysis; for example, regression by cell type used ‘cell type’ as the *Main Covariate*, while analyses that further stratified by mutation type (i.e., C>T, C>G, etc.) additionally included ‘mutation type’ as a *Secondary Covariate*.

We then used the emmeans package^80^ to obtain from models the estimated mutation rates and their 95% confidence intervals for every combination of levels of the *Main Covariate* and levels of any specified *Secondary Covariates*, as well pairwise mutation rate comparisons (rate ratios, 95% CIs, and *P-*values) between levels of the *Main Covariate* within each *Secondary Covariate* stratum (if specified). Specifically, we used the ‘emmeans’ function with parameters ‘specs = “*pairwise ∼ Main Covariate* | *Secondary Covariate 1* (if specified) ∗ *Secondary Covariate 2* (if specified) ∗ …’, ‘type = response’, and ‘offset = 0’ followed by the ‘summary’ function with parameters ‘infer = TRUE’, ‘level = 0.95’, ‘adjust = mvt‘, and ‘cross.adjust = sidak‘. The ‘offset = 0’ parameter outputs rates per unit exposure (i.e., mutations/[bp · yr]), the adjust parameter applies the ‘multivariate t’ (mvt) method to adjust for multiplicity of contrasts among levels of *Main Covariate* within each *Secondary Covariate* stratum, and the cross.adjust parameter applies the Sidak method to adjust for multiplicity across strata. For mutation burden versus age plots that show a 95% confidence band for the regression line, the confidence band was obtained via a parametric bootstrap using the fitted model with 500 iterations. Plots were made with the ggplot2 package^75^.

### Mutational signature analysis

Mutational spectra and signature analyses used the set of unique mutations (i.e., if a mutation was detected more than once in a sample, it was counted as a single mutation). Mutational spectra of indels were tabulated with indelwald^81^ and spectra of indels and substitutions were plotted with sigfit^82^. Mutational signature analyses were performed with sigfit^82^. All sigfit functions for signature extraction and fitting utilized the parameters ‘model’ = “multinomial”, ‘iter’ = 10000 (10,000 total iterations), ‘warm’ = 5000 (5,000 warm-up iterations), and ‘control=list(adapt_delta = 0.99)’ (decreases step size during modeling). After each signature extraction, we refit to the extracted signatures to obtain more accurate signature exposure estimates. Reconstruction accuracy was calculated as the cosine similarity of spectra reconstructed based on signatures and their exposures to the original spectra. Per-signature mutation counts and burdens were calculated as [signature’s exposure estimate] × [total mutation count or burden]. Per-signature mutation rates and mutation rate comparisons were obtained using the same generalized linear modeling process described above. COSMIC signatures used during analysis were obtained from v3.4 of the COSMIC catalogue.

#### Substitutions

For substitution analyses of individual samples, we excluded samples from the three individuals who were 2 years old due to their low total mutation counts (range 34 – 180 mutations; average 77 mutations) that yield unreliable mutational spectra. All substitution signature analyses corrected for the difference of the distribution of trinucleotides in interrogated base pairs relative to the genome using the ‘opportunities’ parameter of the sigfit functions. Specifically, this was set to a matrix with the ratio of counts of each trinucleotide context in interrogated base pairs relative to the genome, for each sample. The relatively small size of our sample cohort makes de novo extracted signatures less reliable than COSMIC catalogue signatures that were extracted from thousands of samples^61,83,84^, especially in terms of identifying signatures such as SBS1 that contribute less to the overall spectrum^2^ and in separating signatures that share features likely due to ‘cross-contamination’ such as SBS5 and SBS16 as previously described^46^. Therefore, after performing de novo signature extraction, we fit our data to the COSMIC catalogue signatures that most closely resemble the extracted signatures rather than fitting directly to the de novo extracted signatures.

For the de novo signature extraction phase, we iteratively: (a) extracted signatures (beginning with two signatures using sigfit’s ‘extract_signatures’ function) and then, (b) fit our data to the COSMIC catalogue signatures most closely resembling the extracted signatures while simultaneously extracting one additional signatures using sigfit’s ‘fit_extract_signatures’ function. This process was repeated until the additional extracted signature showed very low similarity any COSMIC signature. The comparison to the COSMIC catalogue used the cosine similarity metric and excluded COSMIC signatures that resemble SBS5 per a cosine similarity analysis we performed among all pairs of COSMIC signatures (SBS3, SBS25, SBS39, SBS40a/b/c, SBS89, SBS92, SBS96), as such signatures can be mis-assigned to samples due to their similarity to SBS5^85^. Applying this approach to spectra of individual samples yielded in the first extraction two signatures that most closely resembled the ubiquitous signatures SBS1 and SBS5 (cosine similarities 0.70 and 0.89, respectively)^2,44,46^. In the second iteration we extracted an additional signature that most closely resembled SBS16 (cosine similarity 0.75).

The next extraction iteration did not yield a signature with significant similarity to a COSMIC signature (maximum cosine similarity = 0.60). To further confirm the robustness of the identified active signatures, we also applied this signature extraction and fitting approach to spectra that combined all mutations from samples of the same cell type stratified the transcription level genome regions (i.e., intergenic, non-expressed genes, and expressed gene regions Q1, Q2, Q3, and Q4; each for PN and GN, yielding 12 total spectra), as transcription is a major driver of differences in PN and GN mutational patterns. In the first extraction step, this yielded two signatures that most closely resembled SBS1 and SBS5 (cosine similarities 0.73 and 0.89, respectively). In the second iteration we extracted an additional signature resembling SBS16 (cosine similarity 0.74), and subsequent extracted signatures did not yield a signature with significant similarity to a COSMIC signature (maximum cosine similarity = 0.61).

#### Indels

Due to the small number of indels per sample, we performed indel signature analysis after pooling samples into four age groups: 2-29 yo (6 individuals), 30-53 yo (5 individuals), 54-79 yo (5 individuals), ≥ 80 yo (5 individuals). De novo extraction yielded indel signatures with low cosine similarity to the COSMIC catalogue except for an ID9-like signature (cosine similarity 0.78) that was identified when extracting two signatures, so we proceed to fit our data using a set of COSMIC indel signatures (ID1, ID2, ID4, ID5, ID8, ID9, and ID11) that are either prevalent across diverse tissues or that were observed in prior somatic mutation studies of neurons^44,46^.

## Extended Data Figure Legends

**Extended Data Figure 1.**
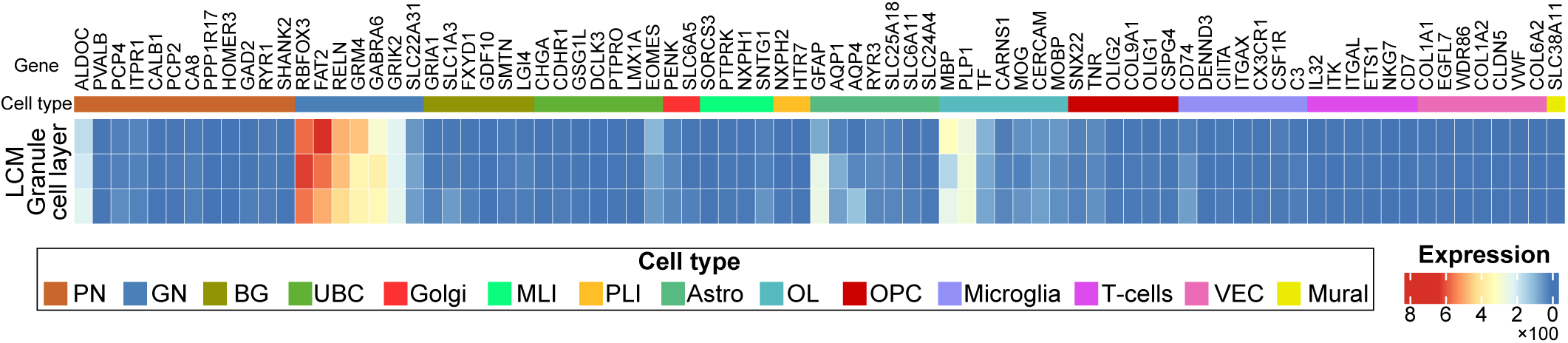
Smart-seq3 bulk RNA-sequencing of granule cell layer laser-capture microdissection. Normalized unique molecular index (UMI) exon-aligned read counts obtained by Smart-seq3 bulk RNA-seq of three laser-capture microdissections from the granule cell layer (**Methods**; top to bottom, individuals 937, 5609, 11589) for marker genes of major cerebellar cell types. See **Supplementary Table 3** for data of all genes. GN: granule neurons; PN: Purkinje neurons; BG: Bergmann glia; UBC: unipolar brush cells; Golgi: Golgi neurons; MLI: molecular layer interneurons; PLI: Purkinje layer interneurons; Astro: astrocytes; OL: oligodendrocytes; OPC: oligodendrocyte progenitor cells; VEC: vascular endothelial cells; Mural: mural cells.

**Extended Data Figure 2.**
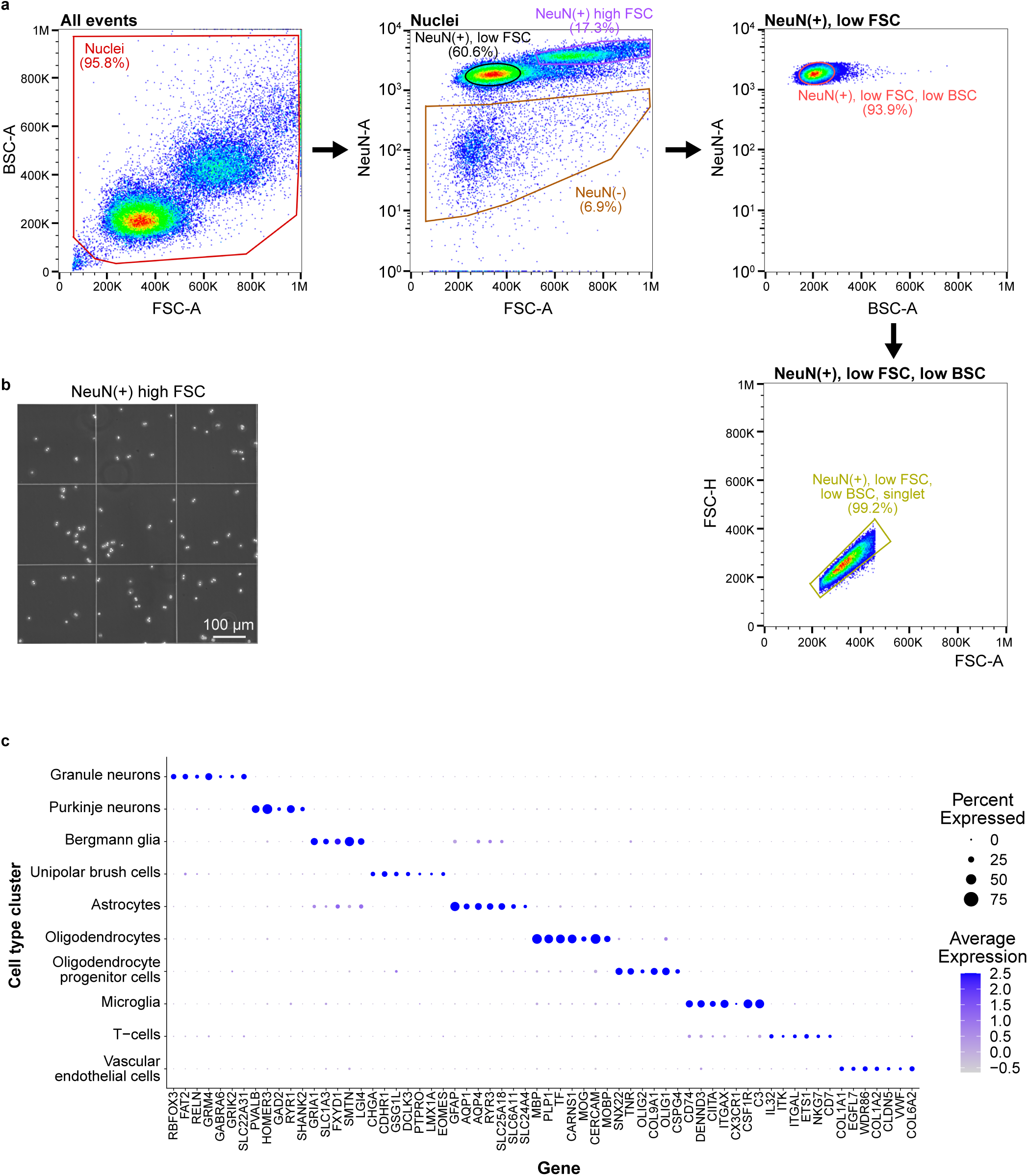
Fluorescence-activated sorting of granule neuron nuclei. **a**, Gates used for fluorescence-activated nuclei sorting. The ‘NeuN(+), low FSC, low BSC, singlet’ gate was used for granule neuron mutation profiling. This gate and the ‘NeuN(-)’ gate were used for snRNA-seq quantification of cell types. Note, FSC and BSC are the two scatter gates available in the sorting instrument. Plots were made with FlowJo software after downsampling to 50,000 randomly selected events from a representative sample from individual 937 that was sorted for snRNA-seq. Annotated gate percentages are the percentage of included events out of the parent gate. A, area; K, thousand; M, million. **b**, phase-contrast microscopy on a hemocytometer of sorted ‘NeuN(+) high FSC’ events, which were not sorted for mutation profiling or snRNA-seq, confirming these are multiplets of nuclei, likely mostly multiple granule neuron nuclei. **c**, snRNA-seq expression of marker genes in each annotated cell type cluster in the UMAP plot in Fig. 1d. See **Methods** for sources used to select these marker genes.

**Extended Data Figure 3.**
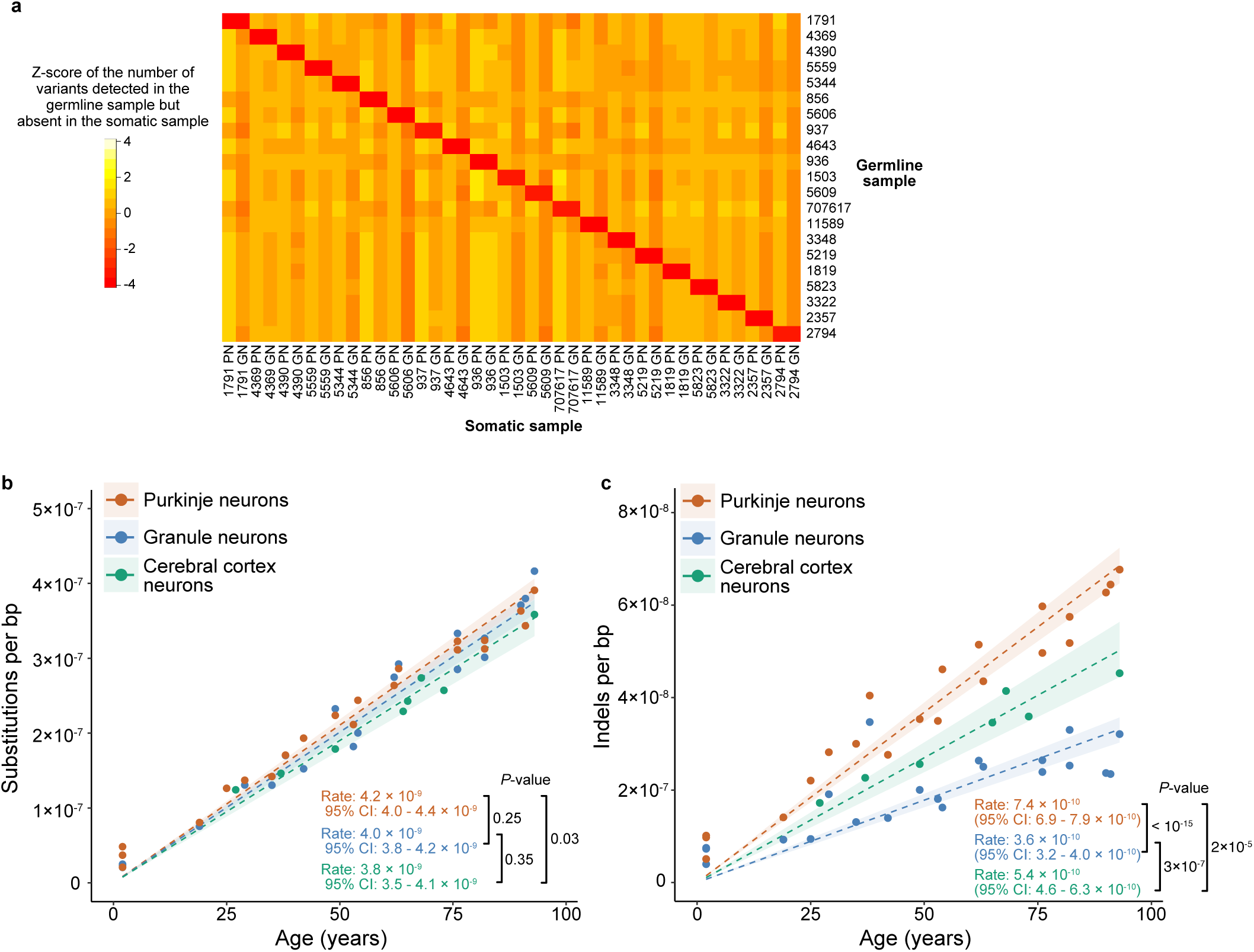
Substitution and insertion/deletion mutation rates. **a**, Fingerprinting of sample identities using germline variants (**Methods**). The z-score is centered and scaled per row (i.e., per germline sample). **b,c**, Substitution (b) and indel (c) mutation burdens versus age, including data from a prior duplex sequencing study of cerebral cortex neurons^2^. Dashed lines show predictions of the regression model (**Methods**). Shaded regions show 95% confidence bands (**Methods**). Rates are in units of mutations/[bp · year]. *P* values are adjusted for multiplicity of tests (**Methods**). See **Supplementary Tables 7,8** for further details of rates and comparison statistics. PN, Purkinje neurons; GN, granule neurons; CI, confidence interval.

**Extended Data Figure 4.**
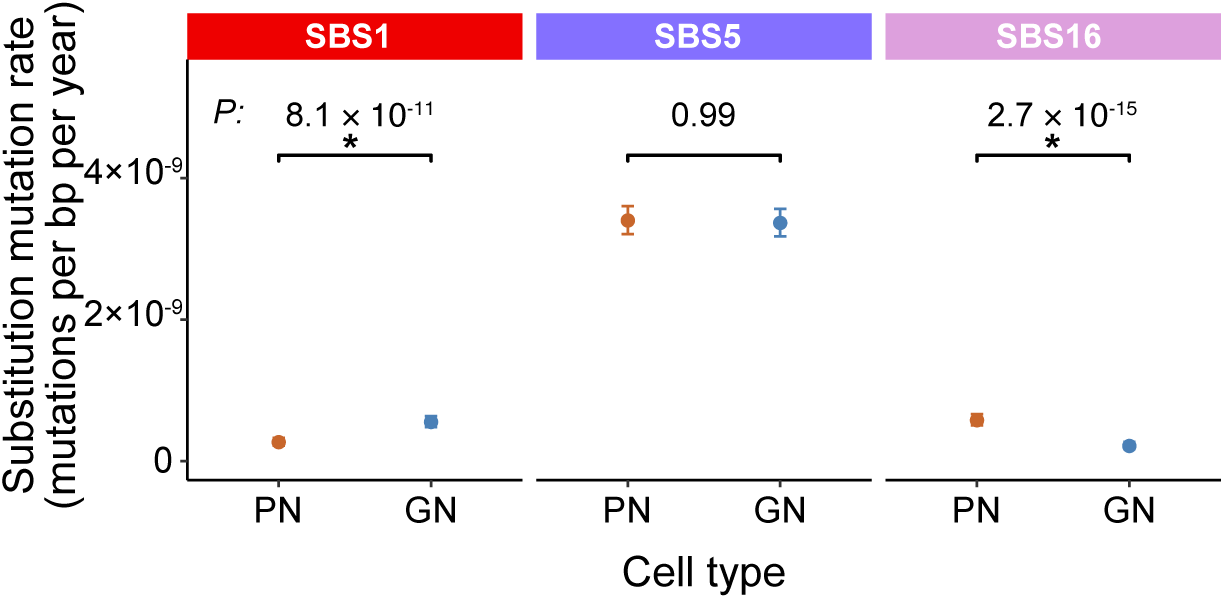
Mutation rates of substitution and signatures. Substitution mutation rates for each signature. PN versus GN comparisons are annotated with *P*-values (asterisks mark significant differences), and error bars show 95% CIs (**Methods**). PN, Purkinje neurons; GN, granule neurons.

**Extended Data Figure 5.**
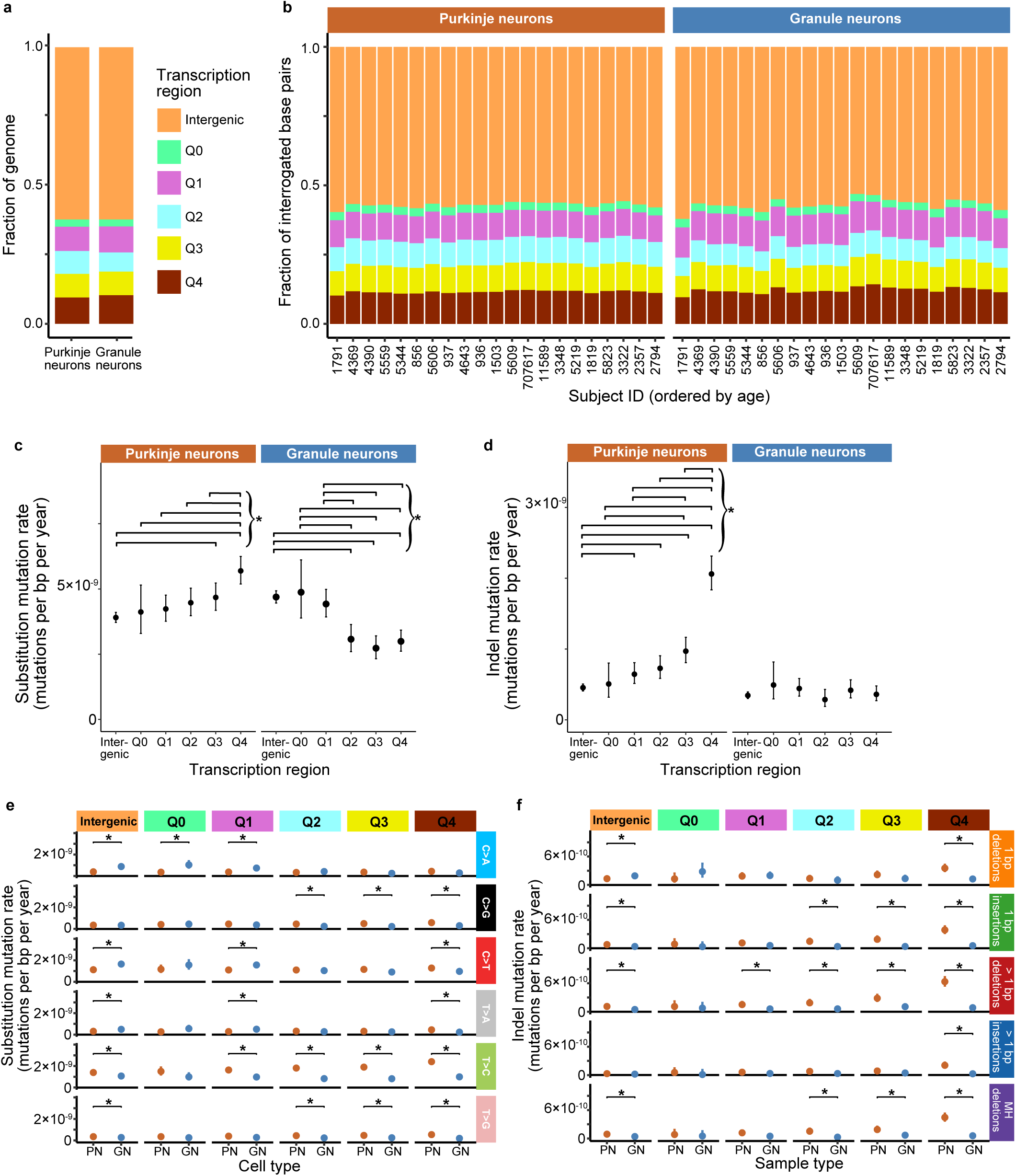
Associations with transcriptional activity. **a,b,** Fraction of the genome (a) and fraction of interrogated base pairs per sample (b) in each transcriptional activity region set (**Methods**). Q0 is regions of genes that are not expressed. Shared legend is in (a). **c,d,** Comparisons for substitution (c) and indel (d) mutation rates in PNs and GNs between different transcription regions. Error bars show 95% CIs. **e,f,** Comparisons of substitution (e) and indel (f) mutation rates between PNs and GNs in different transcription regions, stratified by mutation type. **c-f,** Statistically significant comparisons out of every possible pairwise comparison (corrected for multiple hypothesis testing) are marked. See **Supplementary Tables 7,8** for further details of rates and comparison statistics, including *P*-values. **f,** MH, microhomology.

**Extended Data Figure 6.**
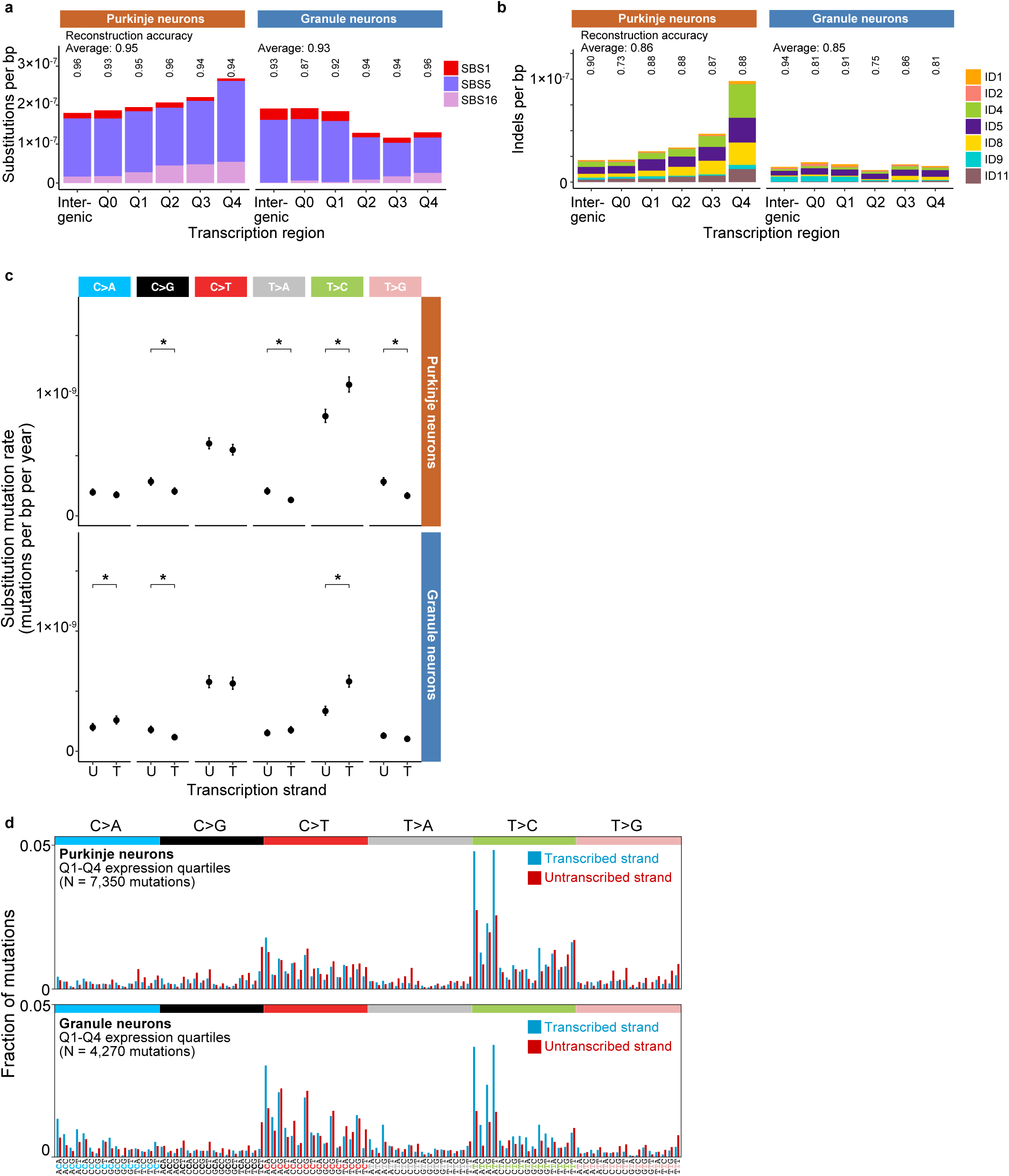
Additional analyses of mutation rates stratified by transcription strand and activity. **a,b,** Burdens of substitution (a) and indel (b) signatures after pooling all samples of each cell type by transcription region. Reconstruction accuracy is annotated above each sample, as well as the average reconstruction accuracy across samples of each cell type. See **Supplementary Table 10** for exposure estimates. **c,** Substitution mutation rates for the untranscribed versus transcribed strand for different mutation types. See **Supplementary Tables 7,8** for further details of rates and comparison statistics, including *P*-values. U, untranscribed strand; T, transcribed strand. **d,** Mutation spectra stratified by strand combined across all samples of each cell type for unique mutations in expressed gene regions. Spectra were corrected for the trinucleotide distribution of interrogated bases relative to the genome.

**Extended Data Figure 7.**
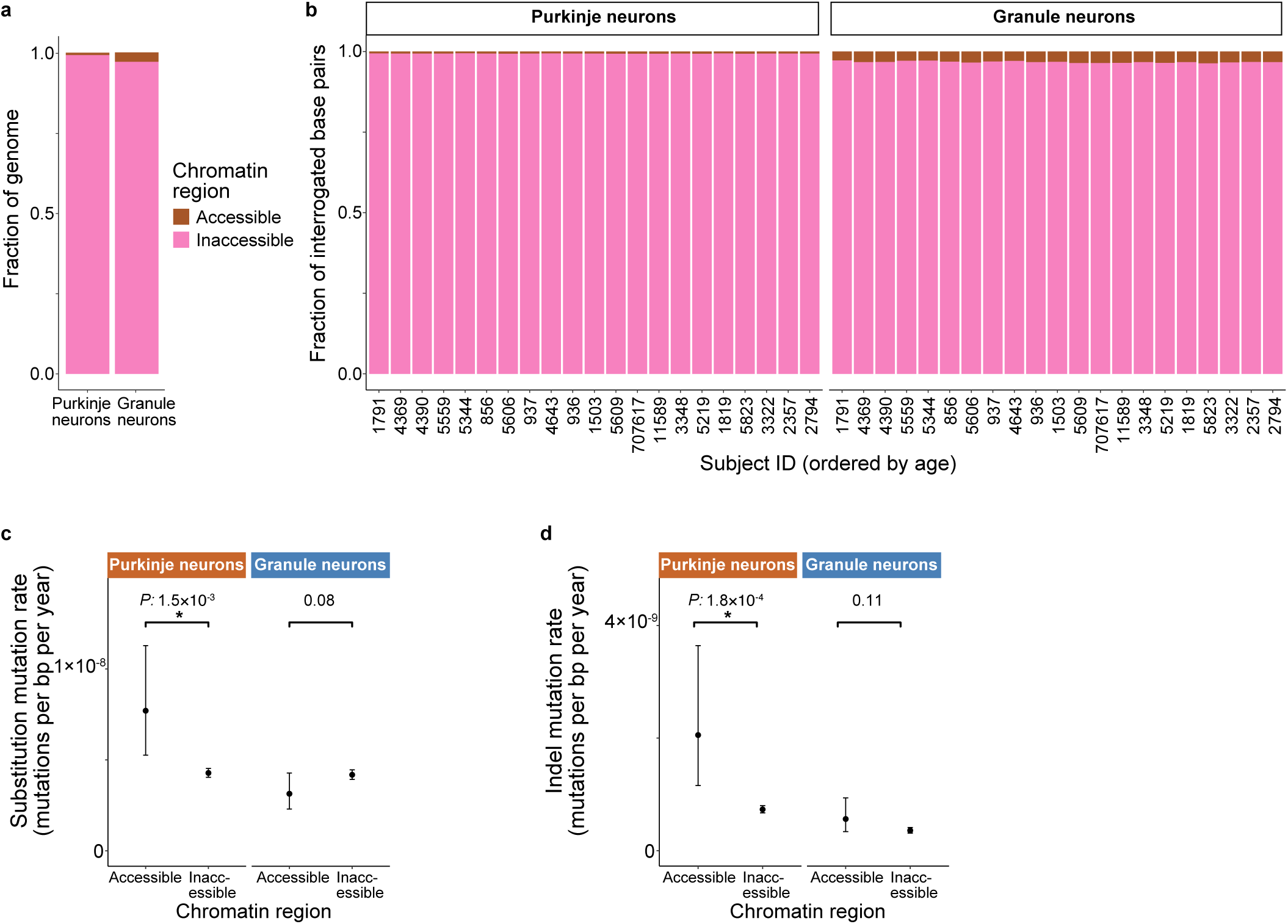
Associations with chromatin accessibility. **a,b,** Fraction of the genome (a) and fraction of interrogated base pairs per sample (b) in each chromatin region (**Methods**). Shared legend is in (a). **c,d,** Comparisons for substitution (c) and indel (d) mutation rates in PNs and GNs between different chromatin regions. Comparisons are annotated with *P*-values (asterisks mark significant differences), and error bars show 95% CIs (**Methods**). See **Supplementary Tables 7,8** for further details of rates and comparison statistics.

## Acknowledgements

This work was supported by grants from the Pew Charitable Trusts (G.D.E.), the National Institute on Aging (T32AG052909 and F32AG076287; M.G.P.), and the NIH Common Fund (UH3NS132024; G.D.E). We thank the NIH NeuroBioBank for providing human tissues. Sequencing performed at the New York University (NYU) Grossman School of Medicine Genome Technology Center was supported in part by the National Cancer Institute (P30CA016087) and a National Institutes of Health Shared Instrumentation Grant (1S10OD023423-01). The computational work was supported in part by the New York University Information Technology High Performance Computing resources, services, and staff expertise, and by the New York University Grossman School of Medicine High Performance Computing Core. We thank the NYU Grossman School of Medicine’s Center for Biospecimen Research and Development core (Tomoe Shiomi and Luis Chiriboga) for assistance with tissue sectioning and the Experimental Pathology Research Laboratory (Cynthia Loomis and Branka Dabovic) for assistance with laser capture microdissection.

## Author Contributions

M.G.P. and G.D.E. conceived the project and designed the experiments. M.G.P. performed the experiments. M.G.P., A.S., and G.D.E. analyzed the data. M.G.P. and G.D.E. wrote the manuscript, with input from A.S.

## Competing Interests

The authors do not have competing interests.

## Supplementary Information

<u>Supplementary Tables.</u> This file contains **Supplementary Tables 1-11**. **Supplementary Table 1** contains details of profiled individuals and samples. **Supplementary Table 2** contains a list of all samples and sequencing metrics. **Supplementary Table 3** contains normalized UMI counts for the Smart-seq3 experiment. **Supplementary Table 4** contains the counts and percentages of each cell type for the snRNA-seq experiment. **Supplementary Table 5** contains substitution and indel mutation burdens. **Supplementary Table 6** contains a list of all detected somatic mutations, with genomic position, reference base(s), and mutant base(s) for each mutation annotated as in VCF (Variant Call Format) files. **Supplementary Table 7** contains statistics of mutation rates. Q0, regions of genes that are not expressed. **Supplementary Table 8** contains statistics of mutation rate comparisons. Q0, regions of genes that are not expressed. PN, Purkinje neurons; GN, granule neurons; CN, cerebral cortex neurons. **Supplementary Table 9** contains mutational spectra for substitutions and indels for unique mutations (i.e., counting only once those mutations that are detected more than once within a sample). It also contains the ratio of the fraction of base pairs with each trinucleotide context in the final interrogated base pairs relative to the fraction of base pairs with that trinucleotide context in the reference genome, which was used to correct the number of observed mutations for differences in trinucleotide distributions between the final interrogated base pairs and the genome (**Methods**). **Supplementary Table 10** contains the exposure estimates for each sample to substitution and indel mutational signatures. Q0, regions of genes that are not expressed. **Supplementary Table 11** contains a list of regions used for gene expression and chromatin accessibility regions after merging overlapping regions within each region set.

## Data availability

Sequencing data generated in this study are available at the NCBI Sequence Read Archive (Accession ID pending). See **Supplementary Table 2** for accession IDs of each sample.

## Code availability

The NanoSeqTools package^45^ is available at https://github.com/evronylab/NanoSeqTools.

## Notes

### Competing Interest Statement

The authors have declared no competing interest.

